# Single-cell analysis of early chick hypothalamic development reveals that hypothalamic cells are induced from prethalamic-like progenitors

**DOI:** 10.1101/2021.04.09.438683

**Authors:** Dong Won Kim, Elsie Place, Kavitha Chinnaiya, Elizabeth Manning, Changyu Sun, Weina Dai, Kyoji Ohyama, Sarah Burbridge, Marysia Placzek, Seth Blackshaw

## Abstract

The hypothalamus is an evolutionarily ancient brain region that regulates many innate behaviors, but its development is still poorly understood. To identify molecular mechanisms controlling hypothalamic specification and patterning, we used single-cell RNA-Seq to profile multiple stages of early hypothalamic development in the chick. We observe that hypothalamic neuroepithelial cells are initially induced from prethalamic-like cells. Two distinct hypothalamic progenitor populations emerge later, which give rise to paraventricular/mammillary and tuberal hypothalamus. At later developmental stages, the regional organization of the chick and mouse hypothalamus closely resembles one another. This study identifies selective markers for major subdivisions of the developing chick hypothalamus and many uncharacterized candidate regulators of hypothalamic patterning and neurogenesis. As proof of concept for the power of the dataset, we demonstrate that follistatin, a novel prethalamic progenitor-like marker, inhibits hypothalamic induction. This study both clarifies the organization of the early developing hypothalamus and identifies novel molecular mechanisms controlling hypothalamic induction, regionalization, and neurogenesis.

**Highlights:**

- Early hypothalamic development was profiled in chick using scRNA-Seq and multiplexed HCR.
- Hypothalamic cells are induced from prethalamic-like neuroepithelial cells.
- Distinct paraventricular/mammillary and tuberal progenitor populations emerge later, and hypothalamic organization is evolutionarily conserved.
- Prethalamic progenitor-derived follistatin inhibits hypothalamic specification.

**Graphical Abstract:** 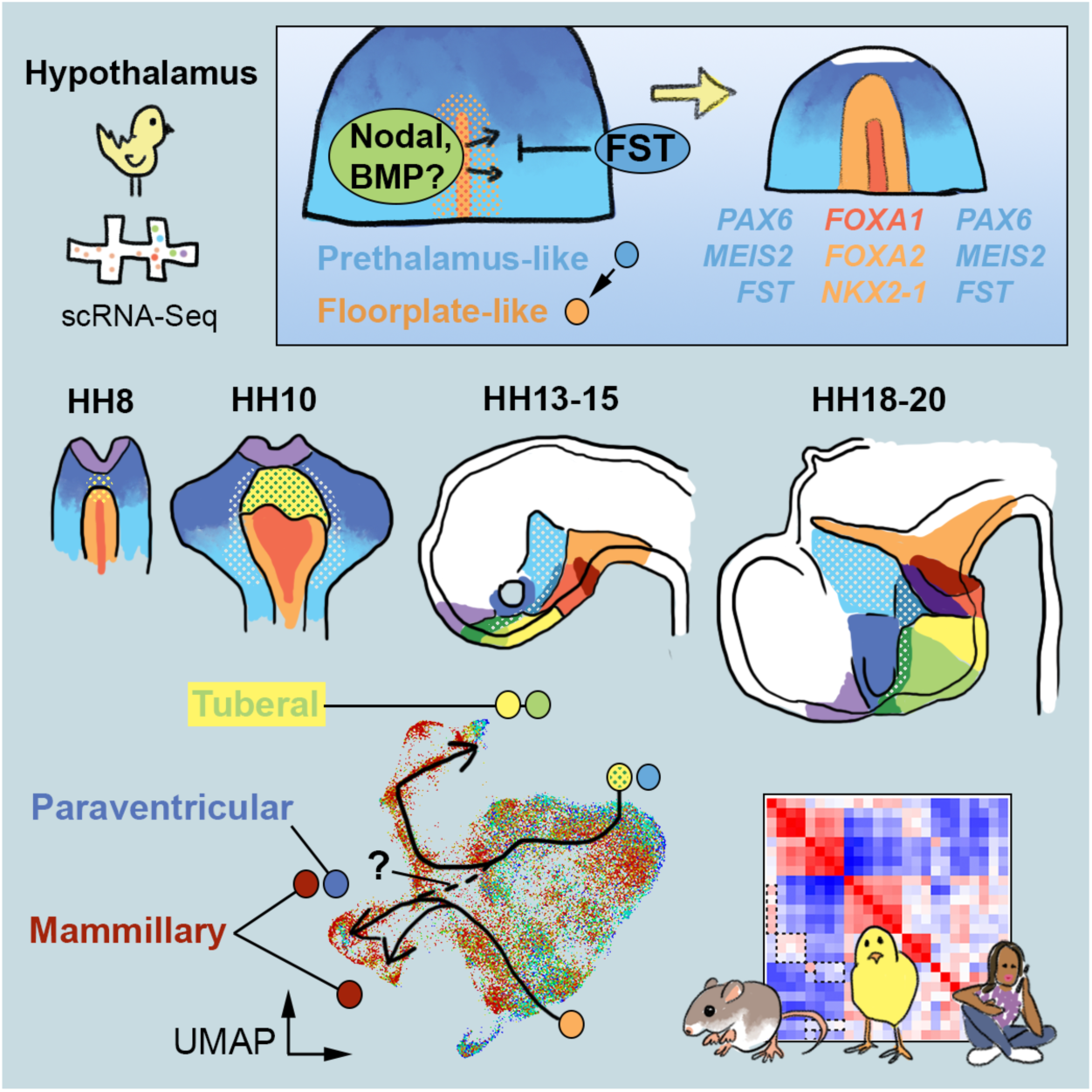

## Introduction

The hypothalamus is a central regulator of a broad range of behaviors and homeostatic physiological processes. These include control of circadian rhythms, the sleep-wake cycle, hunger and thirst, response to stress, and a range of social, sexual, affiliative, and emotional behaviors (Saper and Lowell, 2014; Swaab, 2003). The increased prevalence of homeostatic metabolic disorders such as obesity and diabetes, as well as psychiatric disorders such as depression, underline the importance of a better understanding of hypothalamic organization and dysfunction. Studies of early hypothalamic development can provide important insights into the overall structure and function of this highly complex brain region. Moreover, a better characterization of the molecular pathways that direct hypothalamic development is key to understanding how disruption of hypothalamic development can lead to metabolic and behavioral disorders in adulthood (Bedont et al., 2015; Biran et al., 2015; Xie and Dorsky, 2017).

Recent years have seen substantial progress in our understanding of the molecular mechanisms controlling hypothalamic neurogenesis. Building on previous work that identified molecular markers of major hypothalamic regions (Shimogori et al., 2010), several studies have used single-cell RNA sequencing (scRNA-Seq) analysis (Kim et al., 2020; Lee et al., 2018; Romanov et al., 2020) to comprehensively profile dynamic changes in gene expression that correlate with the specification of major hypothalamic neuronal subtypes. In combination with traditional genetic studies (Aslanpour et al., 2020; Bedont et al., 2014; Chen et al., 2020; Kim et al., 2020, 2021; Liu et al., 2015; Lu et al., 2013; Romanov et al., 2020; Salvatierra et al., 2014), scRNA-Seq has now begun to identify key molecular regulators of the development of major hypothalamic neuronal and glial cell types in mice. However, these studies have not yet provided insight into the earliest phases of hypothalamic development, corresponding to the period in which neuroepithelial cells first acquire hypothalamic identity and become regionalized. This is largely because of the extremely small size of the preneurogenic mouse hypothalamus, and the difficulty in obtaining defined developmental stages, and has left fundamental questions about the organization of the early hypothalamus unanswered.

The relatively large chick embryo, in which early hypothalamic development occurs over a considerable period, offers an attractive alternative system for addressing these questions (Blackshaw et al., 2010; Burbridge et al., 2016). A range of classical embryological and pharmacological manipulations, coupled with fate mapping, have defined the precise period during which hypothalamic cell identity is induced in chick, identified a number of robust molecular markers of the early hypothalamus that are broadly evolutionarily conserved, and identified key extrinsic signals that control early hypothalamic patterning, including Nodal, SHH, BMP2/7, and FGF10 (Burbridge et al., 2016; Dale et al., 1997; Fu et al., 2017; Placzek and Briscoe, 2005). However, due to the high morphological complexity and dynamic gene expression profile of the developing chick hypothalamus, it has been difficult to directly link these early developmental findings with the later processes of hypothalamic regionalization and neurogenesis, which have been more extensively studied in mice. The advent of new technologies such as scRNA-Seq and multiplexed hybridization chain reaction (HCR) provide powerful tools to address this knowledge gap.

In this study, we used scRNA-Seq to comprehensively profile changes in gene expression in the chick hypothalamus across six different time points, covering the full period between hypothalamic induction and the onset of regionalization and neurogenesis. This, in combination with multiplex HCR analysis, allowed us to define the major spatial domains at each of these stages. We find that at Hamburger-Hamilton (HH)8, the earliest stage profiled, the hypothalamus exists as floor plate-like neuroepithelial midline cells, characterized, at their anterior limit, by a cap of *BMP2/DBX1*/*NKX2-1*-positive cells, and located immediately adjacent to prethalamic-like progenitors. Using RNA velocity to infer cell lineage relationships (La Manno et al., 2018), we show the emergence of hypothalamic from prethalamic-like clusters. We describe how early hypothalamic neuroepithelial cells undergo a dramatic expansion, giving rise to progenitors that are already becoming molecularly and spatially organized into mammillary/paraventricular and tuberal subsets by HH13. By HH18-20, we observe a pattern of expression of region-specific markers that closely reflects patterns previously reported in neurogenic mouse and human hypothalamus. As proof of principle for the power of this analysis to identify novel mechanisms controlling hypothalamic patterning, we tested the effects of gain and loss of function of follistatin, which we find to be selectively expressed in prethalamic-like progenitor cells. *In vivo* and *ex vivo* analysis revealed that follistatin promotes prethalamic-like identity while inhibiting hypothalamic identity, demonstrating an unexpected role for prethalamic-derived signals in controlling early hypothalamic development. This study provides a molecular roadmap for early hypothalamic development at unprecedented spatial and temporal resolution, confirms the extensive evolutionary conservation of the hypothalamus, reveals unexpected molecular homologies among developing hypothalamic regions, and identifies new signaling pathways controlling hypothalamic induction and regionalization.

## Results

### ScRNA-Seq analysis of early chick hypothalamic development

To comprehensively examine gene expression across early chick hypothalamic development we profiled six-time points corresponding to the period of hypothalamic induction (HH8), hypothalamic patterning and regionalization (HH10, HH13/14, and HH15/16), and the initiation of neurogenesis (HH13/14, HH15/16, HH18/19, and HH20/21) (Fig. 1A)(Dale et al., 1997, 1999; Ware et al., 2014, 2016). At each stage, neural tissue was isolated, and morphological landmarks were used to isolate hypothalamic tissue from surrounding brain areas (Fig. 1B). Between 4,000 and 20,000 cells were analyzed at each stage, with 76,000 cells analyzed in total (Fig.S1). UMAP datasets were created for each stage (Fig. 1C). Initial interrogation revealed minimal contamination from *FOXG1+* telencephalic or *PITX3-*positive mesencephalic cells at any age (Fig. 1D; Table ST1). Small clusters, separate from the main body, and detected between HH8 and HH13 (Fig. 1C,D), represented contaminating cells from underlying tissues. *PAX6/MEIS2-*positive prethalamic-like progenitor cells (see below) were detected at every age and were the predominant cell type detected at HH8 (Fig. 1D). At all later stages, however, the great majority of all cells profiled were of unambiguous hypothalamic identity (Fig. 1D).

**Figure 1.**
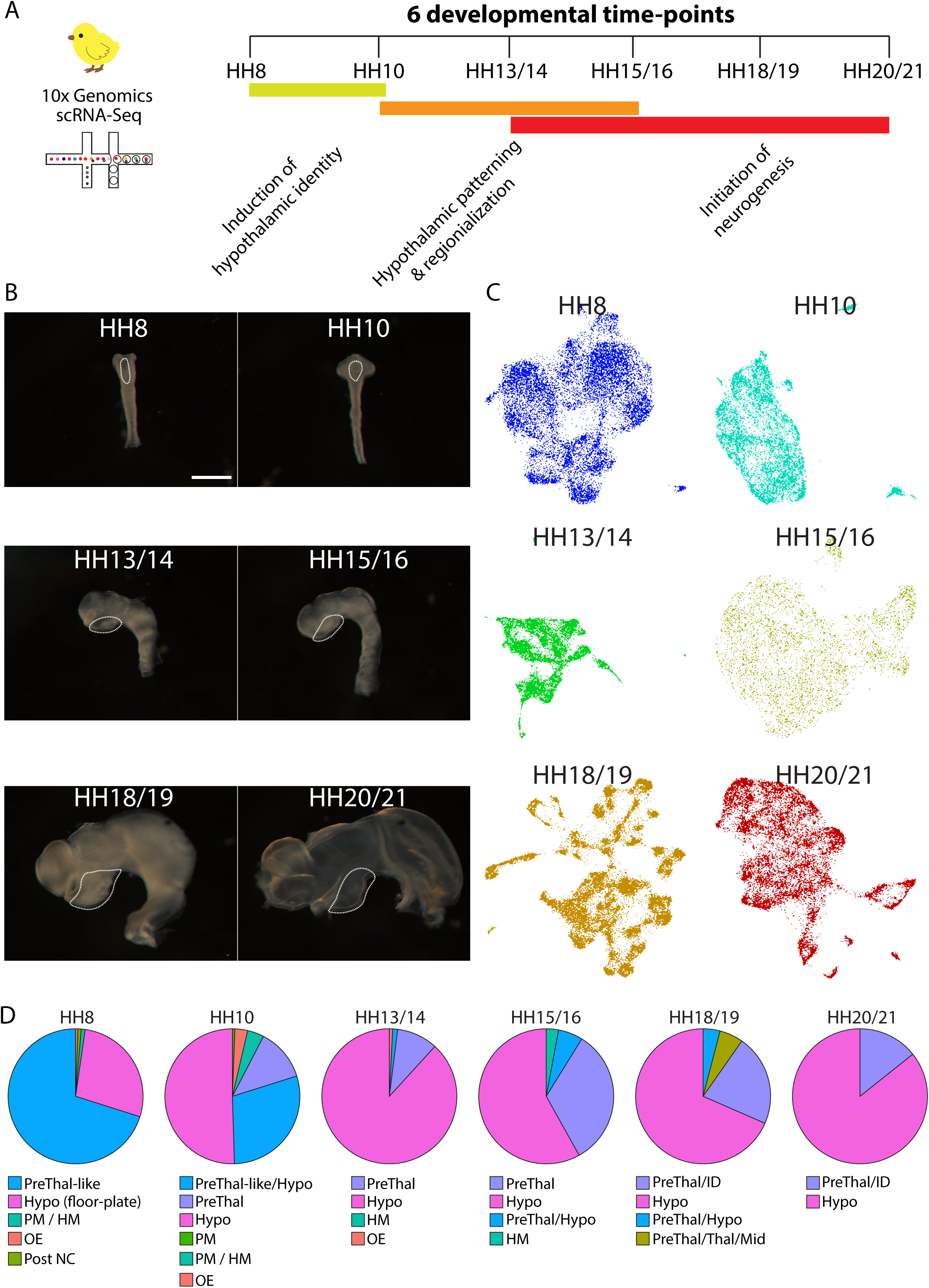
Overview of the generation of the chicken developmental hypothalamus scRNA-Seq dataset. (A) Schematic diagram showing scRNA-Seq experimental design. Hypothalamic tissue was isolated at six developmental time-points (HH8, HH10, HH13/14, HH15/16, HH18/19, and HH20/21) that cover the full course of hypothalamic induction, patterning, regionalization, and initiation of neurogenesis. (B-C) Wholemount views (B) of isolated neuroectoderm (all to same scale), showing dissected areas (white dotted regions), and UMAP plots (C) at HH8, HH10, HH13/14, HH15/16, HH18/19, HH20/21. (D) Pie charts showing the distribution of prethalamic/hypothalamic progenitor cells and contaminating tissues across the 6 developmental time-points. Scale bar = 800 μm. HM = Head mesoderm/mesenchyme, Hypo = Hypothalamus, ID= Intrahypothalamic diagonal, Mid = Midbrain, OE = Oral ectoderm, PM = Prechordal mesoderm, Post NC = Posterior neural crest, PreThal-like = Prethalamus-like progenitors, PreThal = Prethalamus, Thal = Thalamus.

### Specification of hypothalamic identity

The HH8 hypothalamic scRNA-Seq dataset showed a clear division into two major cluster types: floor plate-like clusters (C1-C3, C5) defined through the expression of *FOXA1/2* and *SHH* (Fig. 2A) (Dale et al., 1999) and prethalamic-like clusters (C8-C10), defined through expression of markers, previously characterized in chick (*PAX6*, *DLX2* (Fig. 2A) (Ferran et al., 2008; Larsen et al., 2001) or mouse (*OLIG2*, *MEIS2, SP8*) (Kim et al., 2020; Shimogori et al., 2010) (Fig. 2A). Floor plate-like cells separated into further distinct clusters based on the expression of molecular markers, including *SST*, *OLFML3*, *CHRD*, *NOV (CCN3),* and *CA2,* many of these not previously described in the neural tube midline (Table ST1). Multiplexed HCR analysis demonstrated the expression of such genes in intermingled midline domains along the anteroposterior axis (Fig. 2C-2E; Fig.S2A-G,O-Z,B,D’-E’,G’). This analysis also identified previously uncharacterized molecular markers of the prethalamic-like region, including *FGF18,* and *FST*. HCR analysis confirmed that *PAX6*, *MEIS2,* and *FST* expression overlapped in a region of the anterior-lateral neural tube that will give rise to the prethalamus and paraventricular nucleus (Fig.S2E-G;Y,A’,C’) (Cobos et al., 2001). Importantly, three of the four floor plate-like clusters (C1, C2, and C3) expressed both floor plate-like markers (*SHH, FOXA2*), a subset of prethalamic-like markers (*PAX6, MEIS2*, *FST*), and, additionally, well-characterized markers of hypothalamic cell types (*NKX-2.1*, *POMC*, *BMP2, DBX1, SIX6*) (Table ST1). HCR analysis revealed expression of these genes in a horseshoe-like cap wrapping around floor plate-like midline cells, intermingling with them at their anterior-most limit (Fig. 2C, 2E; Fig.S2B,G). These, and additional HCR analyses (Fig.S2), suggest the location of emerging hypothalamic cells at HH8 (Fig. 2I).

**Figure 2.**
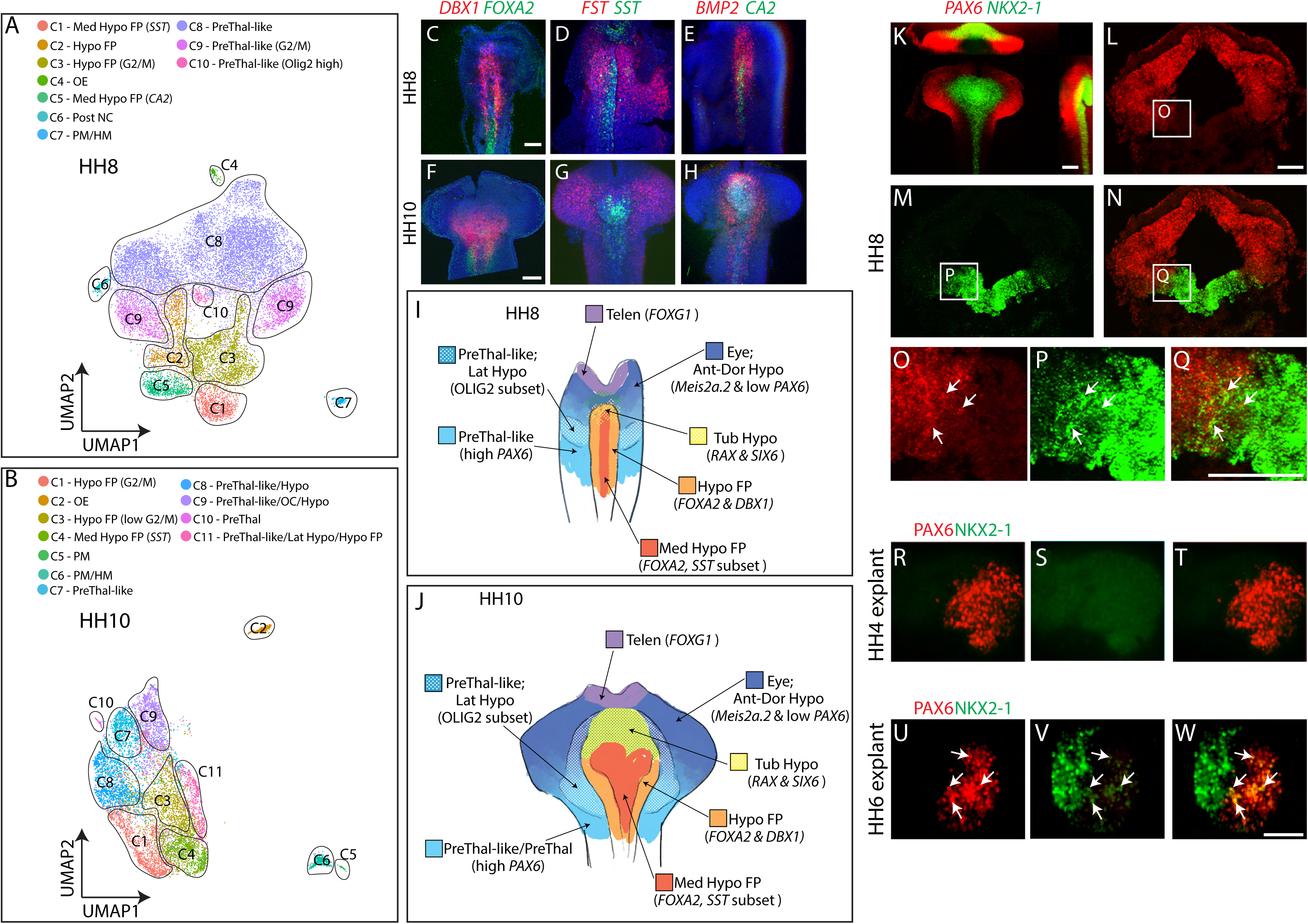
Specification of hypothalamic identity between HH8 and HH10. (A) UMAP plot showing the distribution of prethalamic-like and hypothalamic floor plate-like cells at HH8. (B) UMAP plot showing the distribution of prethalamic-like and hypothalamic cells at HH10. (C-H) Maximum intensity projections after wholemount HCR *in situ* hybridization on isolated neuroectoderm showing *DBX1*/*FOXA2* (C, F), *FST*/*SST* (D, G), *BMP2/CA2* (E, H) at HH8 (C-E), and HH10 (F-H). (I-J) Schematic diagrams showing prethalamic and hypothalamic regions at HH8 (I) and HH10 (J). (K-Q) HCR *in situ* hybridization of *Pax6* and *NKX2-1* at HH8. (R-W) Immunohistochemical analysis of PAX6 and NKX2-1 in HH4 cultured explant (R-T), HH6 cultured explant (U-W) (n=5 explants/condition). Scale bar = 100 μm. Ant = Anterior, Dor = Dorsal, FP = Floor plate, HM = Head mesoderm, Hypo = Hypothalamus, Lat = Lateral, Med = medial, NC = Neural Crest, OE = Oral ectoderm, PM = Prechordal mesoderm, PreThal-like = Prethalamus-like progenitors, PreThal = Prethalamus, Post = Posterior, Telen = Telencephalon, Tub = Tuberal.

At HH10, a broadly similar organization of clusters is apparent: four clusters (C1, C3, C4, and C11) are dominated by *SHH-*expressing cells, and three clusters (C7, C8, and C9) by cells expressing prethalamic-like genes (*PAX6*, *MEIS2, FST*) (Fig. 2B) (Table ST2). In *SHH-*expressing clusters, hypothalamic genes (*NKX2-1, NKX2-4*) that were detected weakly and in sparse numbers of cells at HH8 are now expressed strongly and in many cells (Table S2). Further, some prethalamic-like clusters (C8, C9) now also contain cells expressing low levels of hypothalamic progenitor markers (*RAX, SIX6, FOXA2, SHH, NKX2-2*)(Table S2). These observations indicate that hypothalamic progenitor cells substantially increase in number between HH8-HH10 (Fig. 1D). In support of this, multiplex HCR analysis reveals a substantial lateral expansion of *DBX1, FOXA2, SHH*, *NKX2-1,* and *CA2* (Fig. 2F, 2H, and S2H,J-LN), and an expanded concentric arc of *BMP2* expression (Fig. 2H). *DBX1* and *SIX6*, whose expression overlapped at HH8, now begin to resolve into discrete spatial domains: *SIX6* expression becomes pronounced around the anterior midline (Fig.S2I) (likely prefiguring future anterior ventral regions, including optic midline and tuberal hypothalamus) and *DBX1* becomes more posteriorly-confined (Fig. 2F). *BMP7* and *SST* are nested within the *DBX1*-expressing domain and show a more modest lateral expansion (Fig. 2F, 2G; Fig.S2H). *FST, SP8,* and *PAX6* expression remain strong in lateral regions (prospective prethalamus and optic vesicle) (Fig. 2G; FigS2M,N; FigS4B,Q). *OLIG2* expression, which at HH8 was starting to become concentrated in the prethalamic region, is now also weakly expressed in a concentric ring (Fig.S2K) resembling the expression domain of *BMP2* (Fig. 2H). Together, these studies indicate the rapidly changing gene expression profile of nascent hypothalamic cells at HH10 (Fig. 2J).

The broadly similar cluster organization seen in these two samples, combined with the major changes in the relative sizes of the clusters expressing these markers, led us to aggregate the HH8 and HH10 samples, which facilitates identification of genes that potentially regulate early stages of hypothalamic patterning (Fig.S3A). Immature prethalamic-like cells (*PAX6, DLX2, SP8, MEIS2, FST*-positive) were organized into three distinct clusters (all represented by C5) (Fig.S3A,B) that were partly distinguished through the expression of sex chromosome-specific genes such as *WPKCI-7* (Fig.S3B). A somewhat more mature prethalamic-like cluster (C6) expressed higher levels of *OLIG2* (Fig.S3B) and *HES5* (Fig.S3C). A cluster of floor plate-like hypothalamic cells (defined through *SHH, CHRD*, *CA2,* and *SST* expression) was evident (C1), as were three clusters consisting of *SHH/NKX2-1/NKX2-4*-positive hypothalamic progenitor cells (C2/C3/C4). Of these, cells in cluster C2 expressed relatively higher levels of *NKX2-2, OLIG2, VAX1, HES5,* and *SIX6*, and only low levels of *OLFML3* and *CHRD*, consistent with this cluster being composed of cells of more anterior and lateral (including future tuberal, antero-tuberal and paraventricular) hypothalamic identity. Clusters C3 and C4 closely resembled one another and were largely distinguished through differential expression of cell cycle regulators (in C4,, Table ST3). C3 and C4 expressed low levels of *CHRD* and the secreted Wnt antagonist *SFRP2,* and similar levels of *FOXA2* to those seen in the C1 hypothalamic floor plate-like cluster (Fig.S3B,C, Table ST3).

Despite the similarities between the clusters at HH8 and HH10, we nonetheless observe major changes in their composition (Fig.S3D) in keeping with the observation that hypothalamic progenitor cells significantly expand over this period. RNA velocity analyses, both at individual time-points (HH8 and HH10) and on the integrated HH8/HH10 dataset revealed trajectories from prethalamic clusters to the emerging/nascent hypothalamic clusters (*NKX2-1*/*NKX2-4*-positive) (Fig.S3E-G).

Pseudotime analysis demonstrates that cells on these differentiation trajectories downregulate many prethalamic markers, including *FST*, *DLX2*, and *FGF18*, and upregulate hypothalamic markers such as *NKX-2.1* and *NKX2.4*, potentially indicating that hypothalamic cells arise from prethalamic-like cells (Fig.S3H). Although prethalamic markers such as *PAX6* and *SP8* are also upregulated in the pseudotime analysis, we reasoned that this might simply reflect the presence of maturing prethalamic cells. To directly ask if hypothalamic genes are upregulated in prethalamic-like cells during this period, we performed HCR analyses at HH8-10. This suggests the presence of cells co-expressing *NKX-2.1* and *PAX6* located lateral to the midline (Fig. 2K-Q), and in posterolateral regions, *SST*/*SP8-*, *NKX2-1*/*SP8*- and *NKX2-1/DBX1*-positive cells (Fig.S4). This implies that prethalamic-like cells may undergo conversion to a hypothalamic identity during this period. Previously, we have explanted and cultured prospective hypothalamic cells at neural plate stages of development, to show that hypothalamic specification, as measured by *NKX2-1* expression, has not yet begun at HH4, but has been initiated by HH6 (Ohyama et al., 2005). Using this assay we found evidence for the direct conversion of prethalamic-like to hypothalamic identity: in cultured HH4 explants, the majority of cells contained PAX6 protein, but no NKX2-1 was detected, while in HH6 explants, NKX2-1-positive cells were detected, many of which co-expressed PAX6 (Fig. 2R-W).

RNA velocity analysis of the integrated (HH8/HH10) dataset revealed a second, clear differentiation trajectory that connects hypothalamic floor plate-like cells (C1) to cells that resemble posterior hypothalamic (including future mammillary) progenitors (C3/C4) and then to hypothalamic cells (C2) (Fig.S3I). In this trajectory, cells lose expression of markers such as *SST* and *CHRD* relatively early, show a more gradual decline in *FOXA2* expression, while progressively upregulating hypothalamic progenitor markers such as *NKX-2.1*, *NKX-2.2,* and *NKX-2.4* and the mammillary progenitor marker *PITX2* (Fig.S3I). Taken together, this suggests the existence of two distinct developmental trajectories for generating hypothalamic progenitors.

### The emergence of hypothalamic regionalization and initiation of neurogenesis

The scRNA-Seq datasets obtained from HH13 and HH15 showed broadly similar clusters (Fig.S5, Table ST4,ST5), and were aggregated for analysis (Fig. 3A). By these stages, the organization of the hypothalamus has radically changed, with regional identity emerging and the first neurons being generated. As at HH8 and HH10, we observe three sub-regions containing mainly intrahypothalamic diagonal (ID) and prethalamic progenitors (collectively termed C4: identified through the expression of *OLIG2, PAX6, SP8, EMX2*), which are distinguished from each other by expression of sex chromosomal genes, and higher expression of *GBX2* and *ZIC2* in the outermost cluster (Fig. 3B,C, Table ST6). RNA velocity analysis indicates that cells in the innermost part of cluster C4 give rise to cells in the outermost part of the cluster (Fig. 3D). The remaining clusters are hypothalamic, distinguished by *NKX2-1, NKX2-4,* and weak *NKX2-2* expression (Fig. 3B). While no postmitotic prethalamic neuronal precursors are observed at this stage, we observe two separate hypothalamic neurogenic trajectories. RNA velocity analysis indicates that the first branch arises from a progenitor cluster (C3) distinguished by the expression of floor plate-like markers such as *FOXA1/2*, *OLFML3*, and *CHRD*. Cells in cluster C3, and the neurogenic clusters C1 and C2, express posterior hypothalamic markers such as *PITX2* and *NKX6-2* (Fig. 3B,C). Adjacent to cluster C3 is another progenitor cluster (C5), distinguished by *SST* expression (Fig. 3C). RNA velocity analysis further reveals that a subset of C5 progenitors gives rise to C3 progenitors (Fig. 3D).

**Figure 3.**
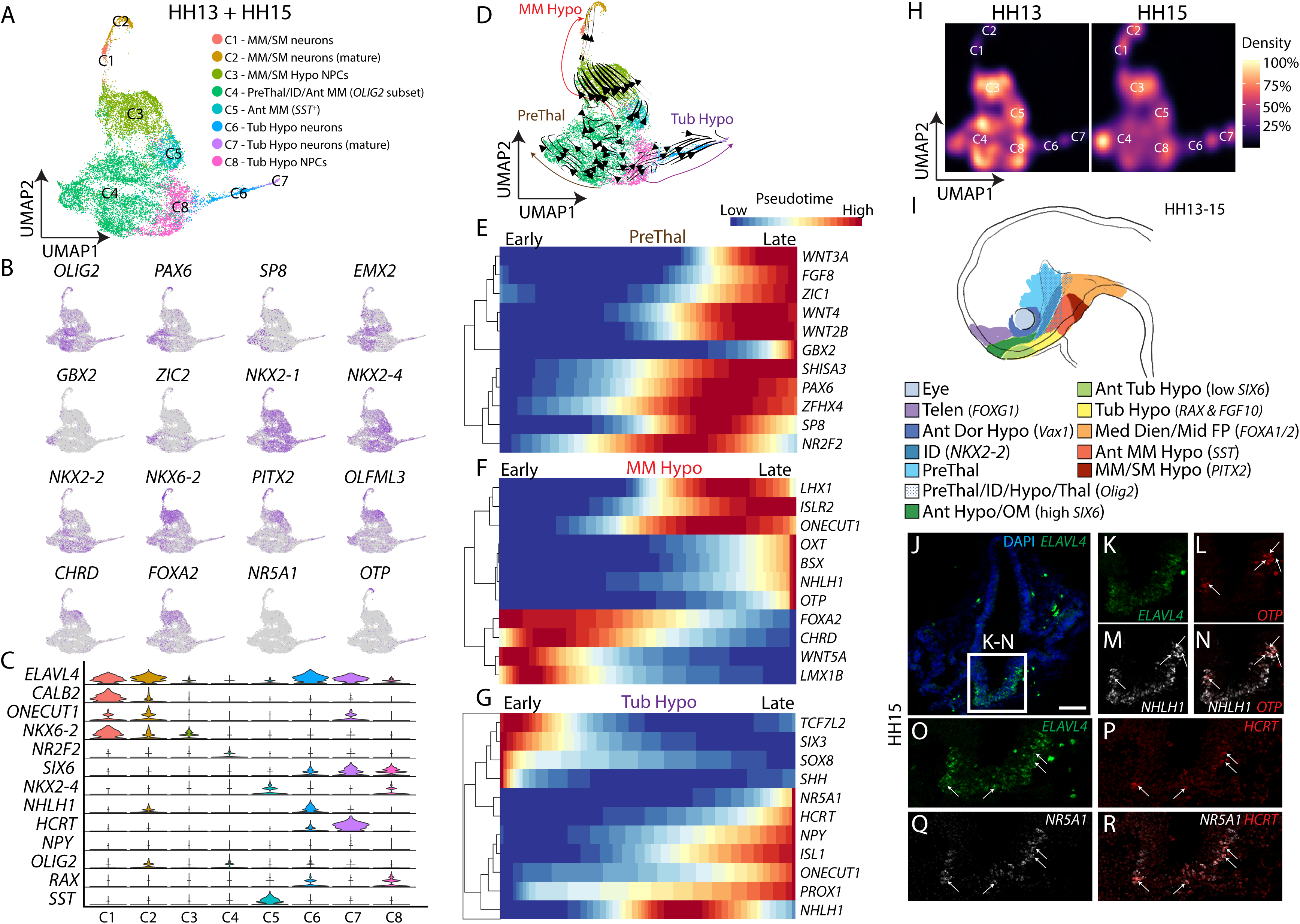
Regionalization of the hypothalamus and initiation of neurogenesis. (A) UMAP plot showing the distribution of hypothalamic regionalization at HH13/HH15. (B) UMAP plot of HH13/ HH15 showing hypothalamic regionalization genes. (C) Violin plot of HH13/HH15 showing hypothalamic regionalization and neurogenesis genes. (D) UMAP plot showing HH13/HH15 scRNA-Seq with prethalamus, tuberal hypothalamus, and mammillary hypothalamus trajectories obtained from RNA velocity. (E-G) Pseudotime analysis of prethalamus (E), mammillary hypothalamus (F), and tuberal hypothalamus (G). (H) UMAP plot showing the distribution of HH13 and HH15 cells. (I) Schematic diagram of HH13/HH15. (J-R) HCR *in situ* hybridization showing a section through the tuberal hypothalamus and optic stalk stained for *ELAVL4*, *OTP,* and *NHLH1* (J-N), and *ELAVL4*, *HCRT*, and *NR5A1* (O-R). Scale bar = 100 μm. Markers shown in (K-R) have been digitally overlaid. Ant = Anterior, Dien = Diencephalon, Dor = Dorsal, FP = Floor plate, Hypo = Hypothalamus, ID = intrahypothalamic diagonal, Med = Medial, MM = Mammillary, NPCs = Neural precursor cells, OM = Optic midline, PreThal = Prethalamus, SM = Supramammillary, Telen = Telencephalon, Thal = Thalamus, Tub = Tuberal.

Cells in the neurogenic trajectory that arises from C3 express high levels of the transcription factor *SIM1* (expressed in both paraventricular nucleus and posterior hypothalamus in the chick) (Caqueret et al., 2005) and the calcium-binding protein *CALB2* (Fig 3C, Table ST6), which is selectively expressed in developing and mature neurons in the mammillary and paraventricular nuclei in mice (Shimogori et al., 2010). Cells at the distal tip of this trajectory strongly express the transcription factors *NHLH1* (Fig. 3C) and *OTP* (Fig. 3B), as well as lower levels of neurohormones *AVP* and *OXT* (Table ST6), all of which are later selectively expressed in the paraventricular nucleus (Xu et al., 2020). This implies that neuronal precursors in this trajectory may give rise to both mammillary and paraventricular neurons.

RNA velocity analysis reveals that a second hypothalamic neurogenic differentiation trajectory arises from a progenitor cluster (C8) which expresses high levels of *SIX6*, *RAX,* and *FGF10* (Fig. 3C; Table ST6). Based on previous work, this is likely to include tuberal hypothalamic progenitors (Fu et al., 2017; Kim et al., 2020; Shimogori et al., 2010). In keeping with this, the neural precursors arising from C8 express genes such as *NR5A1* (Fig. 3B), as well as *NPY* (Fig. 3C) and *POMC* (Table ST6), which selectively mark the ventromedial and arcuate nuclei respectively, as well as genes such as *ISL1* (Table ST6), expressed broadly in mouse tuberal hypothalamic neurons (Shimogori et al., 2010). Unexpectedly, this also included the arousal-promoting neuropeptide *HCRT* (Fig. 3C), which is selectively expressed in the lateral hypothalamus in mice, where it is not detected until the postnatal period. Furthermore, while *LHX9* is selectively expressed in *HCRT*-positive neurons, and is required for *HCRT* expression in both mice and zebrafish (Dalal et al., 2013; Liu et al., 2015), *LHX9* expression is not detected in C7 *HCRT* cells (Table ST6), marking a dramatic difference in species-specific expression of this key regulator of arousal and sleep. The functional significance of this is unclear.

While strongly indicative of a tuberal differentiation route, we note low levels of *OTP* in clusters C6 and C7, while cluster C8 expresses high levels of *VAX1* (Table ST6), predominantly detected in the paraventricular domain at later stages (see below). Thus, we cannot rule out that cluster C8 includes progenitors that give rise to paraventricular neurons.

UMAP plots at HH13 and HH15 confirm a relative increase in the fraction of cells in neurogenic clusters (C1,2,6,7) and a relative decrease in hypothalamic progenitor cell clusters (C3,5,6,8) over HH13 to HH15 (Fig. 3H). Likewise, pseudotime analysis indicates that cellular expression levels of prethalamic markers such as *PAX6*, *GBX2*, and *ZIC1* progressively increase as these cells mature (Fig. 3E). In the paraventricular/mammillary neurogenic population (C1 and C2), there is a progressive decrease in the expression of floor plate-like markers such as *FOXA2* and *CHRD*, with steadily increased expression of the mammillary markers *LHX1, ONECUT1, and OTP* (Fig. 3F) (Marion et al., 2005; Shimogori et al., 2010). Finally, in the tuberal neurogenic population, we observe a rapid reduction in expression of *SIX3*, *SOX8,* and *SHH*, along with an upregulation of the premammillary-enriched homeodomain factor *PROX1* (Kim et al., 2020; Shimogori et al., 2010), and a transient expression of *NHLH1* (Fig. 3G).

Multiplex HCR analysis on sagittal and transverse sections reveals that the clusters map to spatially-resolving progenitor domains at HH13-HH15 (Fig.S6A-Y, schematized in Fig. 3I). The prethalamus (*EMX2/OLIG2*-positive) lies dorsal and posterior to the hypothalamus (*NKX2-1/SHH*-positive). The *NKX2-2*-positive ID can be distinguished, and also expresses *EMX2* and *OLIG2* in its posterior portion. Within the hypothalamus, *DBX1* and *SIX6* begin to distinguish mammillary/posterior and tuberal/anterior progenitors respectively, weakly overlapping in the tuberal domain (Fig.S6C,D). Anterior-most *SIX6-*positive cells are marked by *NKX2-2* and are adjacent to the *FOXG1-*positive telencephalon (Fig.S6B,D,K). *PITX2* and *FOXA2* overlap with but extend posterior to the *DBX1* domain (Fig.S6C,L,N). *SST* is expressed immediately anterior to *PITX2* and *FOXA2* (Fig.S6L,N,Y), most strongly lateral to the midline, and *OLIG2* and *EMX2* are starting to be weakly expressed just lateral to *SST* (Fig.S6T,W,Y), suggesting that these genes are beginning to distinguish different posterior hypothalamic regions. Finally, multiplex HCR analysis confirms the onset of neurogenesis at these stages, in spatially-discrete regions. *ELAVL4* (Fig.S6Z,A’) is detected in mammillary (*PITX2* and *OTP*-positive) progenitor domain(s), and anterior-ventral progenitor domain(s), and in the latter, co-expressed with *NR5A1* and *ISL1* (Fig. 3J-N;Fig.S6Z-O’). *HCRT* expression is restricted to a subset of *NR5A1*-positive cells (Fig. 3O-R).

### Finalizing the overall plan of hypothalamic organization

The data obtained at HH13/15 suggest that the basic outlines of hypothalamic regionalization have been laid down by this stage. This in turn raises the question of when the overall spatial organization of the developing chick hypothalamus is complete. Analysis of the scRNA-Seq datasets obtained from HH18 and HH20 highlighted numerous molecular markers, in discrete clusters (Fig.S7, Table ST7, ST8). Many of these markers have been previously identified as specifically labeling the major spatial subdivisions of the developing mouse hypothalamus (Shimogori et al., 2010). Using a subset of these markers, we performed multiplex HCR analysis at HH18 and HH20. This revealed that as early as HH18, the regional organization of the chick hypothalamus is remarkably similar to that previously described for the E11.5 mouse (Shimogori et al., 2010) (Fig. 4A). Extending from the posterior border of the telencephalon to the prethalamus, the paraventricular region is characterized by *SIM1, OTP, VAX1,* and *RAX* expression, *SIM1* extending slightly into the *FOXG1*-positive telencephalon (Fig. 4B-F). The *PAX6*/*SP8*/*DLX1/OLIG2*-positive prethalamus is located dorsal and posterior to the paraventricular region; the intrahypothalamic diagonal (ID) and the tuberomammillary terminal (TT), which are fully contiguous and largely molecularly identical with the prethalamus, are readily apparent (Fig.4A, B, O, F’, Table ST7, ST8).

**Figure 4.**
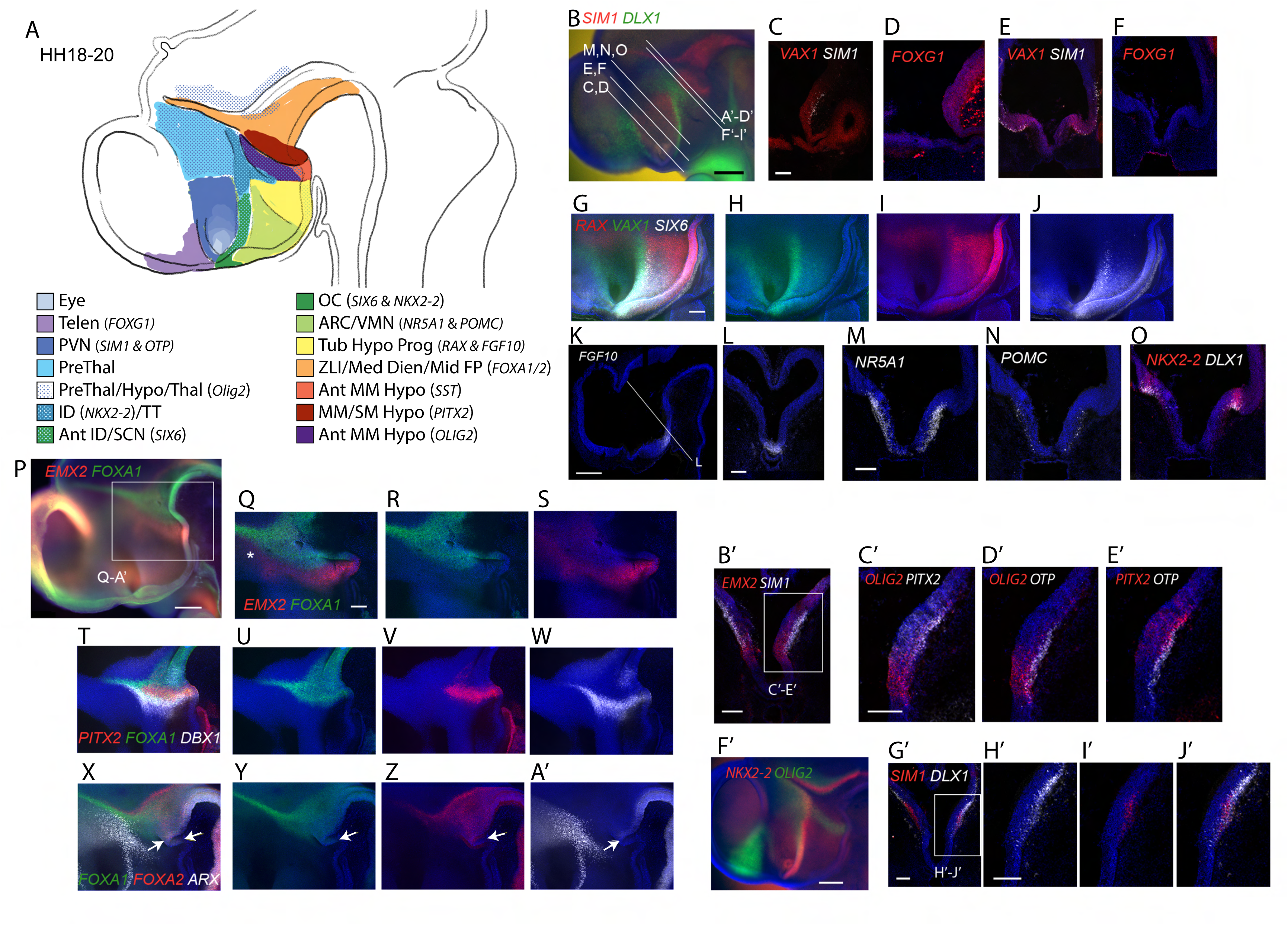
Hypothalamic regionalization at HH18-HH20. (A) Schematic showing major regions of hypothalamus and prethalamus at HH18-HH20, and position of hypothalamic regions relative to Rathke’s pouch. (B) Lateral view of a hemisected embryo after HCR *in situ* to detect *SIM1/DLX1*. (C-F) Transverse sections detecting *VAX1/SIM1* (C, E), *FOXG1* (D, F) at the levels indicated in (B). (G-J) Maximum intensity projections of embryos labeled in wholemount for *RAX/VAX1/SIX6*. (K-O) Transverse (L-O) and sagittal (K) sections detecting *FGF10* (K, L), *NR5A1* (M), *POMC* (N), and *NKX2-2/DLX1* (O) at the levels indicated in (B) and (K). (P) Lateral view of a hemisected embryo showing *EMX2/FOXA1*. Boxed region indicates area shown in (Q-A’). (Q-A’) Maximum intensity projections of embryos stained in wholemount for *EMX2/FOXA1* (Q-S; same embryo as shown in (P)), *PITX2/FOXA1/DBX1* (T-W), *FOXA1/FOXA2/ARX* (X-A’). (B’-E’) Transverse sections stained for *EMX2/SIM1* (B’) and *OLIG2/PITX2/OTP* (C’-E’) at the level indicated in (B). Boxed region in (B’) indicates area shown in (C’-E’) on an adjacent section. (F’) Lateral view of a hemisected embryo stained for *NKX2-2/OLIG2*. (G’-J’) Transverse sections stained for *SIM1/DLX1* at the level indicated in (B). Embryos are HH20 except G-J (HH19) and T-W (HH18). Scale bars represent 100 μm (C, G, L, M, Q, B’, C’, G’, H’) and 250 μm (B, K, P, F’). Markers are shown in (O), (B’-E’), and (G’-J’) have been digitally overlaid. Arrows in (X-A’) indicate anterior limit of *FOXA1/FOXA2*, and *ARX*at the ventral midline. Asterisk = prethalamic *EMX2*.

At this stage, a large domain corresponding to the developing tuberal hypothalamus lies immediately ventral to the ID and dorsal to Rathke’s pouch (Fig. 4A,G-O). Anterior and posterior subdivisions of this region can be distinguished based on the graded expression of *SIX6* and *RAX*. The posterior subdivision is located immediately dorsal to Rathke’s pouch, and is characterized by high *RAX* and *FGF10* (Fig. 4I,K,L). *SIX6* levels are higher and *RAX* lower in the anterior division (Fig. 4G-J). Previous studies, including fate-mapping work (Fu et al., 2017), indicate that the posterior division is composed of tuberal progenitor cells, which give rise to the anterior division composed of anterotuberal progenitors that are likely to give rise to neurons of the ventromedial, arcuate, and anteroventral hypothalamus. At this stage, molecular markers of ventromedial and arcuate nuclei, such as *NR5A1* and *POMC*, are spatially intermingled and their border is not yet defined (Fig. 4M, N). Premammillary-enriched genes such as *PROX1* are also enriched in tuberal-like clusters (Fig.S7, Table ST7, ST8), but a discrete premammillary cluster cannot be distinguished from the broader pool of tuberal hypothalamic progenitors.

Markers that previously defined the prethalamus (*EMX2, OLIG2*) are, at this stage, upregulated in hypothalamic progenitor subsets, and distinct regions within the posterior hypothalamus can at this point be clearly distinguished based on differential expression of *EMX2*, *OLIG2*, *SIM1*, *PITX2*, *FOXA1*, and *FOXA2* (Fig. 4A, examples shown in Fig. 4P-A’,F’). A region expressing *FOXA1*, *PITX2*, low *FOXA2*, and low *EMX2* likely corresponds to the mammillary and supramammillary regions (Fig. 4A). Anterior to this is a narrow band of high *EMX2* expression, which we refer to as anterior mammillary. This zone expresses *OLIG2* and *SST* within a small medial portion (Fig. 4A). *DBX1* appears at the interface between the *EMX2*-/*OLIG2*-positive and *FOXA1*-/*PITX2*-positive regions, partially overlapping both. *SIM1/OTP*-positive neuronal precursors are found in both *EMX2-/OLIG2*- and *FOXA1*-/*PITX2*-positive regions (Fig. 4A,B’-E’). By this stage, *EMX2* and *OLIG2* expression domains extend contiguously into the posterior prethalamus, as seen in the E11.5 mouse (Fig. 4A, P,Q,S,F’) (Shimogori et al., 2010). The intermingling of *SIM1* and *DLX1*-positive cells is observed between the anterior mammillary and dorsal TT regions (Fig. 4B,G’-J’). The posterior limit of the prethalamus and hypothalamus are defined by the *SHH-*positive ZLI dorsally, and ventrally by *ARX*, which marks the di/mesencephalic floorplate (Fig. 4X,A’, arrows), prethalamus, ID, and TT, as in mouse (Fig. 4X,A’) (Shimogori et al., 2010).

### Gene networks controlling hypothalamic regionalization

Since hypothalamic regionalization is largely complete by HH18, we next sought to determine whether scRNA-Seq analysis could help identify gene networks that control this process. Since our samples did not fully cover the full-time course of prethalamic neurogenesis, we removed maturing prethalamic cells (*PAX6/OLIG2/FST*-positive but *RAX/SIX6*-negative), along with the small number of other non-hypothalamic cell types. We then computationally aggregated all scRNA-Seq data from HH8, HH10, HH13, HH15, HH18, and HH20 using UMAP analysis. This revealed a continuous distribution across the time points, which could then be interrogated using both RNA velocity and pseudotime analysis. From HH8 onwards we observe two major populations of progenitors (Fig. 5A,B, Table ST9), corresponding to posterior/mammillary (C0) and tuberal/prethalamic-like (C1) progenitor clusters. RNA velocity plots indicate that both progenitor cell clusters (C0 and C1) converge into a cell cluster (C4) that is distinguished by expression of *ASCL1*, *GADD45G*, and *DLL1*, but which still expresses cell cycle markers such as *CCND1* and *UBE2C* (Fig. 5A,B, Table ST9). Similar populations have been previously observed in scRNA-Seq analysis in developing retina, cortex, and cerebellum, and appear to correspond to progenitors undergoing terminal neurogenic divisions (Carter et al., 2018; Clark et al., 2019; Loo et al., 2019). From this neurogenic progenitor population arise two neuronal differentiation trajectories, corresponding to mammillary/paraventricular (C3) and anterotuberal(C6) cell types (Fig. 5A,B). While C4 is common to both C0 and C1, C1-C6 and C0-C3 trajectories pass through upper and lower subdivisions of C4, respectively. This adds weight to the idea that some paraventricular neurons arise from mammillary progenitors (Fig. 3). We could not confidently identify a second paraventricular trajectory arising from C1, despite HCR showing *SIM1*-expressing cells in close association with *VAX1/RAX*-expressing progenitors (Fig. 4B, C, E, G-I). The fraction of cells in the non-neurogenic progenitor clusters (C0 and C1) remains relatively unchanged from HH10 through to HH18, and decreases only slightly at HH20 (Fig. 5C), indicating that this represents a relatively early stage of hypothalamic neurogenesis, as implied by HCR analysis (Fig. 4).

**Figure 5.**
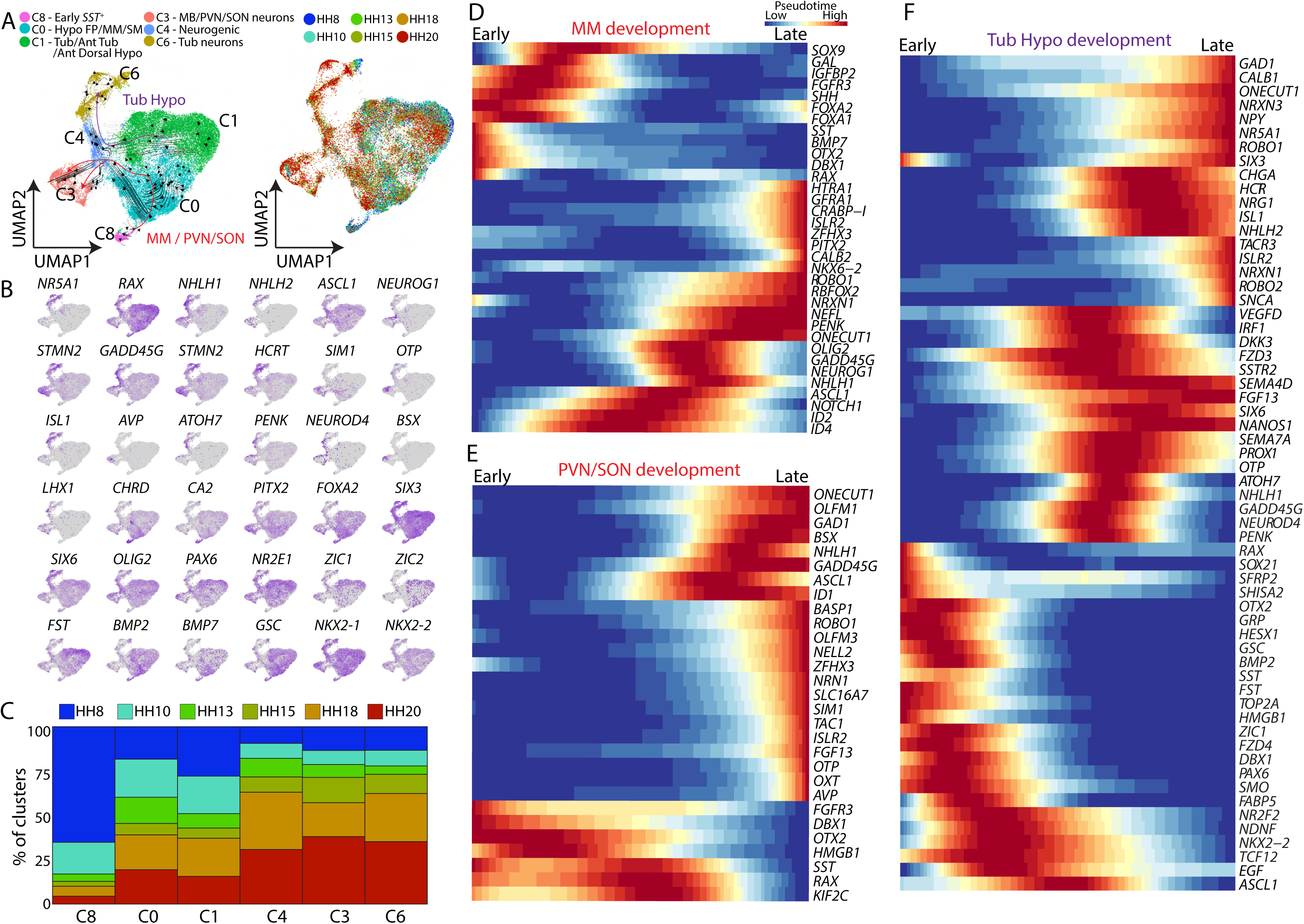
Gene networks controlling hypothalamic regionalization. (A) UMAP plot of clusters across the entire period of hypothalamic development showing the developmental trajectories obtained from RNA velocity analysis of the tuberal hypothalamus, or PVN/SON and mammillary hypothalamus (left), and UMAP plot showing developmental time-points across the entire period of hypothalamic development (right). (B) UMAP plots showing gene expression changes through the period of hypothalamic development. (C) Bar plot showing the distribution of hypothalamic clusters across developmental time-points. (D-F) Pseudotime showing gene expression changes through mammillary (D), PVN/SON (E), and tuberal (F) hypothalamic development. FP = Floor-plate, MM = mammillary, PVN = paraventricular nucleus of the hypothalamus, SON = Supraoptic nucleus, Tub = Tuberal.

Pseudotime analysis identified common and divergent patterns for dynamic gene expression in both neurogenic trajectories. In each, *RAX, DBX1, SST,* and *OTX2* expression are rapidly downregulated (Fig. 5D,E). Posterior/mammillary trajectories initially show persistent expression of floor plate-like cell markers such as *FOXA1/2* and *CHRD*, along with *SHH* and *BMP7*, which are later downregulated. Tuberal/anterotuberal trajectories transiently upregulate prethalamic markers such as *PAX6, ZIC1* and *FST*, but rapidly downregulate these. *BMP2* and *GSC* are initially highly expressed but are also downregulated early on. Neurogenic progenitors in both trajectories selectively express *ASCL1*, *NHLH1*, and *GADD45G*; the paraventricular/mammillary trajectory upregulates *NEUROG1* and *OLIG2*; and the tuberal trajectory upregulates *ATOH7* (Fig. 5F). *ATOH7* has been previously reported to be exclusively expressed in the retina, where it plays an essential role in retinal ganglion cell survival (Brodie-Kommit et al., 2021). This observation suggests *ATOH7* may also play an important role in regulating tuberal hypothalamic development.

Many well-established regional and cell type-specific markers are only strongly detected in neural precursors near the tips of the trajectories, including *NR5A1*, *NPY,* and *HCRT* in the tuberal stream; and *SIM1*, *OTP*, *PITX2*, and *AVP* in the posterior stream (Fig. 5B), consistent with these being upregulated in maturing hypothalamic cells. Both the posterior and tuberal stream bifurcate by HH18 (Fig. 5A). In the posterior trajectory, posterior hypothalamic markers such as *PITX2* and *FOXA2* are distributed across both branches at HH20, but paraventricular nucleus (PVN)-specific transcripts such as *OTP*, *AVP*, and *BSX* are restricted to one branch (Fig. 5B). One interpretation of this observation is that PVN-specific characteristics develop from a program common to mammillary and PVN-like neurons.

### Evolutionary conservation of hypothalamic neuronal precursor identity

To obtain a better understanding of hypothalamic neuronal precursor diversity and evolutionary conservation, we performed a UMAP analysis of the aggregated HH18 and HH20 scRNA-Seq datasets, computationally extracted all postmitotic neural precursors, and then separately clustered these to identify cell type-specific markers (Fig. 6A, Table ST10). We identify eight main clusters, consisting of three tuberal, three mammillary, one paraventricular, and one prethalamic/ID/TT-like cluster. Tuberal clusters (C0, C1, C3) all express *SIX3*, *SIX6*, and *ISL1* (Fig. 6C), but can be distinguished on the basis of other markers. C0 shows high *POMC*, low *HCRT*, *NR5A1*, *NR2F2* expression, and no expression of *NPY* or *NKX-2.1*. C1 shows high *NPY*, *NHLH1*, and *HCRT* expression, low *NR5A1*, *NR2F2*, and *NKX-2.1* expression, and no *POMC* expression. C3 shows high *POMC* and *HCRT* expression, low *NPY* and *NKX-2.1*, and low *NR2F2* expression (Fig. 6B,C). This implies that distinct populations of *POMC* and *NPY*-expressing neurons are already emerging at this age, well in advance of what is seen in mice (Padilla et al., 2010).

**Figure 6.**
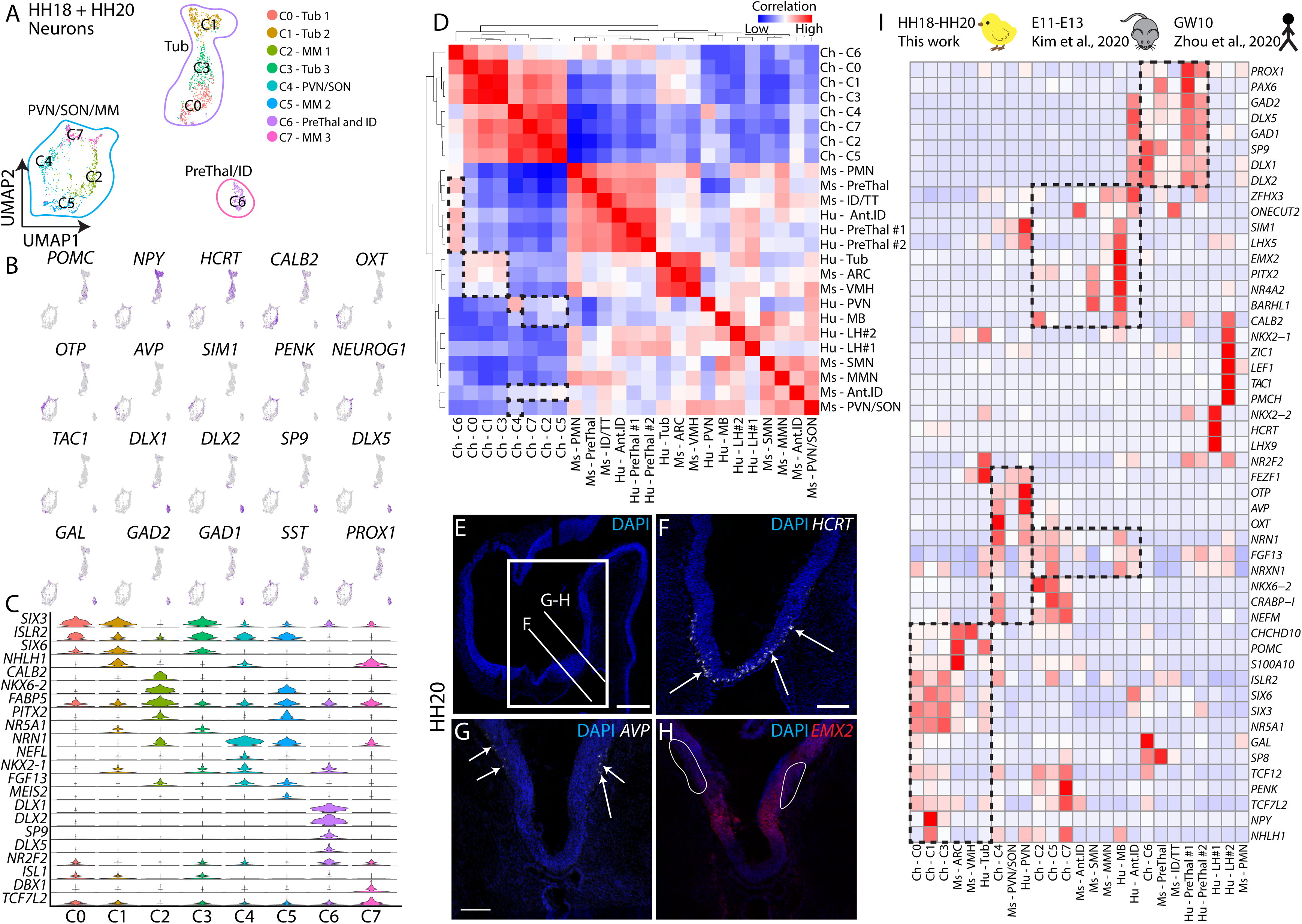
Evolutionary conservation of hypothalamic neuronal precursor identity. (A) UMAP plot showing the distribution of mature hypothalamic and prethalamic/ID neurons at HH18/HH20. (B) UMAP plot showing expression of genes enriched in different hypothalamic neuronal clusters. (C) Violin plot showing hypothalamic cluster-specific genes. (D) Correlation heatmap for developing chicken hypothalamus (HH18/19-HH20/21), mouse hypothalamus (E11-E13, (Kim et al., 2020)), human hypothalamus (GW10, (Zhou et al., 2020)). (E-H) HCR *in situ* hybridization at HH20 showing *HCRT* (F), *AVP* (G), and *EMX2* (H) in transverse sections at the levels indicated in (E). White outline in H indicates location of AVP-expressing cells in F. (I) Heatmap showing conserved key gene expressions between chicken, mouse, and human developing hypothalamus. Scale bar = 100 μm (F-H), 250 μm (E). ID = Intrahypothalamic diagonal, MM = Mammillary, PreThal = Prethalamus, PVN = Paraventricular nucleus of the hypothalamus, SON = Supraoptic nucleus, Tub = Tuberal.

Mammillary (C2, C5, C7) and PVN/SON (C4) clusters are grouped together but can be distinguished by several well-established molecular markers. C4 cells selectively express *SIM1*, *OTP*, *OXT*, *AVP* (Fig. 6B), but also selectively express high levels of *OLIG2* and *NEFL*, and do not express the posterior markers *FOXA1/2* or *PITX2* (Fig. 6C, Table ST10). C2, based on its enriched expression of *SIM1*, *NKX6-2*, and *CALB2* (Fig. 6B), as well as *NKX6-2*, *PITX2*, (Fig. 6C) and *FOXA1/2*, may correspond to neurons of the mammillary nucleus. C5 cells express high levels of *TAC1*, *PITX2, NKX6-2,* and *FOXA1/2* (Fig. 6B, C, Table ST10), and likely correspond to immature supramammillary neurons. C7 cells express high levels of *NKX6-2*, *SST*, *DBX1*, and *OTP*, which at HH20 correspond to part of the anterior mammillary region adjacent to the mammillary (Fig. 6B,C).

GABAergic neurons of the developing hypothalamus are predominantly found in the ID and TT, and are physically contiguous with, and largely molecularly indistinguishable from, neurons of the prethalamus (Kim et al., 2020; Shimogori et al., 2010). Cluster C6 corresponds to these neurons: it selectively expresses canonical markers of hypothalamic GABAergic neural precursors, such as *DLX1/2* and *GAD1/2* (Fig. 6B,C), along with transcription factors such as *SP8* and *SP9* (Table ST10), which are expressed in both prethalamus and more dorsal regions of the ID and TT in mouse (Kim et al., 2020; Shimogori et al., 2010), and which are also found in the chick ID at HH20 (e.g. Fig. 4O). Notably, this cluster shows very few *GSX2*-expressing cells. Since *GSX2* is one of the few prethalamic markers that are not also expressed in the developing hypothalamus (Kim et al., 2020), this indicates that the overwhelming majority of the neurons in this cluster likely represent hypothalamic neurons of the ID and TT.

The availability of scRNA-Seq from early stages of mouse and human hypothalamic neurogenesis makes it possible to directly compare these chick neuronal precursor clusters to clusters previously annotated in E11-E13 mouse (Kim et al., 2020) and gestational week 10 human hypothalamus (Zhou et al., 2020). Tuberal clusters (C0, C1, and C3) closely matched mouse arcuate and ventromedial hypothalamic clusters, as well as the human tuberal hypothalamic cluster (Fig. 6D,I). A few notable differences in expression were, however, observed. As previously mentioned, *NPY* and *NHLH1* expression were highly enriched in chick C1, but not enriched in human or mouse samples. The human tuberal hypothalamus showed much higher relative expression levels of *NR2F2*, *FEZF1*, and *NKX2-1*. Both mouse arcuate and human tuberal clusters also showed substantially higher levels of *POMC* than the chick, while mouse arcuate and ventromedial clusters showed much higher expression of *CHCHD10* than either chick or human tuberal clusters (Fig. 6I).

C4 chick cells most closely resemble the PVN/supraoptic nucleus (SON) mouse cluster and the human PVN cluster (Fig. 6I). In general, the chick more closely resembles the human than it does the mouse, with both chick and human showing high and selective expression of *FGF13*, *NRN1*, *OXT*, and *AVP*. Interestingly, while AVP and OXT-positive neuronal precursors are found in the prospective PVN, some are found in the *EMX2*-positive mammillary region, suggesting that these cells might later either migrate from this region or else only transiently express these markers (Fig. 6G,H). C4 cells selectively express a number of genes not enriched in humans or mice, including *ISLR2*, *TCF12*, *NEFM*, and *CRABP1* (Fig. 6I), but do not express the homeodomain transcription factor *LHX5*, which is expressed in both mouse and human.

Mammillary clusters (C2, C5, and C7) resemble the human mammillary cluster, with C2 sharing its high and selective expression of *CALB2*. While both C2 and C5 express *PITX2*, neither closely resembles the mouse supramammillary cluster (Fig. 6I). The human mammillary cluster, however, also expresses high levels of supramammillary markers such as *NR4A2* and *BARHL1*, suggesting that this represents both mammillary and supramammillary regions (Kim et al., 2020). No clear counterpart for C7 is detected.

Cluster C6 is grouped with both human and mouse ID clusters, as well as human prethalamic clusters (Fig. 6I). C6 showed much higher levels of expression of *GAL*, relatively higher expression of *DLX1/2*, and lower expression of *DLX5* and *GAD2* relative to both mouse and human. Since *DLX1/2* directly regulates *DLX5* expression and precedes the expression of GABAergic markers, this may simply imply that the chick C6 are less mature than their mammalian counterparts and these may represent timing rather than species-specific differences in gene expression. No clear counterparts to either the premammillary cluster in mice or the lateral hypothalamic clusters in humans are detected in chick, consistent with our finding that these structures are not clearly defined at this age.

### Inhibition of hypothalamic induction by prethalamic-derived follistatin

Previous studies have demonstrated a central role for extrinsic signaling in controlling the patterning and regionalization of the developing hypothalamus, including Shh, Wnt, FGF, BMPs, TGFbeta, and Notch/Delta (Aujla et al., 2013; Corman et al., 2018; Fu et al., 2017, 2019; Kapsimali et al., 2004; Lee et al., 2006; Manning et al., 2006; Mathieu et al., 2002; Newman et al., 2018; Pearson et al., 2011; Shimogori et al., 2010). Supporting and extending this work, our scRNA-Seq studies reveal dynamic expression of such key signaling ligands, and their receptors, between HH8 and HH13, along with other receptor-ligand pairs that have not previously been linked to regulation of hypothalamic development (FigS8A,B).

Earlier work has suggested that *Shh* expression in the hypothalamic midline is induced by a combination of Nodal and Shh signaling derived from prechordal mesoderm (Mathieu et al., 2002; Müller et al., 2000; Patten et al., 2003). Strikingly, scRNA-Seq analysis reveals that prethalamic-like cells express high levels of *FST* at HH8 and HH10 (Fig. 2D,G). Since FST is a potent antagonist of signaling by TGFbeta family members, including Nodal, this suggested that prethalamic-derived FST may actively limit the action of prechordal mesoderm-derived Nodal and limit the lateral extent of hypothalamic induction. Indeed, our RNA velocity and HCR analyses at HH8/10 indicated that a subset of prethalamic-like cells adopts a hypothalamic identity, potentially in response to the inductive effects of prechordal mesoderm-derived Nodal (Fig.S2E,F). To directly test this hypothesis, anti-FST antibodies or recombinant FST were injected into the developing hypothalamic region of HH5/6 embryos, which were then allowed to develop to HH14 (Fig. 7A). Exposure to anti-FST antibodies led to a dramatic increase in the size of the *NKX-2.1/SHH-*positive hypothalamic domain relative to controls (Fig. 7B-D,F-H), and a concomitant reduction in the relative size of the prethalamic *PAX6* domain, although *PAX6* expression in Rathke’s pouch was unaltered (Fig. 7E,I). Conversely, no *SHH/NKX-2.1*-positive cells were observed in embryos treated with recombinant FST (Fig. 7J-L). *SHH* expression was detected along the ventral midline, closely abutting the prethalamic domain of *PAX6* expression, which appeared expanded relative to controls (Fig. 7E,M).

**Figure 7.**
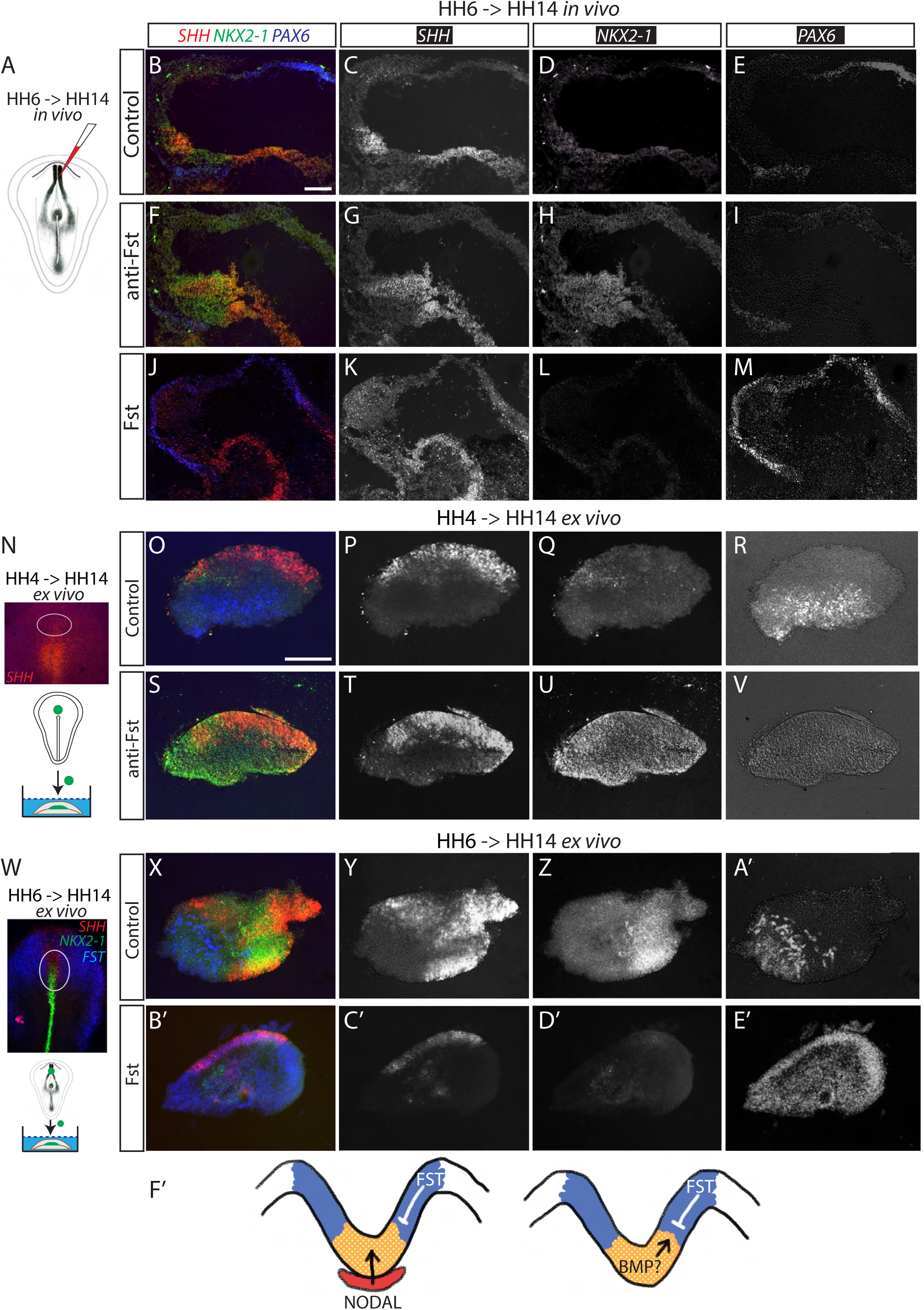
Inhibition of hypothalamic induction by prethalamic-derived follistatin. (A-M) HCR *in situ* hybridization showing *SHH* (B, C, F, G, J, K), *NKX2-1* (B, D, F, H, J, L), and *PAX6* (B, E, F, I, J, M) in sagittal sections of HH14 embryos after *in vivo* injection (A) of PBS (B-E), anti-Fst (F-I), or Fst (J-M) at HH5-6. n=3 embryos/condition. (N-V) HCR *in situ* hybridization showing *SHH* (O, P, S, T), *NKX2-1* (O, Q, S, U), and *PAX6* (O, R, S, V) in sections of explants, cultured from HH4-HH14 (N) in either control medium (O-R), or anti-Fst (S-V). (W-E’) HCR *in situ* hybridization showing *SHH* (X, Y, B’, C’), *NKX2-1* (X, Z, B’, D’), and *PAX6* (X, A’, B’, E’) in sections of explants, cultured from HH6-HH14 (W) in either control medium (X-A’), or anti-Fst (B’-E’). n=7-10 explants/condition. (F’) Schematic showing FST-mediated hypothalamus formation. Scale bar = 100 μm.

In parallel, we performed *ex vivo* studies. Building on previous work that delineated the spatial position of the future hypothalamus and the time at which it is specified (Ohyama et al., 2005), we dissected out prospective hypothalamus at HH4, prior to the age at which it is specified, and removed all traces of prechordal mesoderm. Explants were cultured in 3D collagen gels in either control media or the presence of anti-FST until HH14 equivalent (Fig. 7N). Control explants showed separate domains of *PAX6* and *SHH* expression, with almost no detectable *NKX-2.1* expression (Fig. 7O-R). In contrast, explants exposed to anti-FST downregulated *PAX6* and showed a dramatic upregulation of *NKX-2.1* expression throughout the explant, with many *SHH/NKX-2.1*-positive cells now detected (Fig. 7S-V).

Finally, we dissected out prospective hypothalamus at HH6, when *NKX-2.1* expression is first detected (Fig. 7W). Explants were cultured in control media or the presence of recombinant FST until HH14 equivalent. Robust hypothalamic differentiation was detected in controls, with many *NKX-2.1/SHH*-positive cells detected along with a separate domain of *PAX6*-positive cells (Fig. 7X-A’). In contrast, exposure to FST almost eliminates *NKX-2.1* expression and leads to *PAX6*-positive cells being detected throughout the explant (Fig. 7B’-E’). We conclude that prethalamic-derived FST inhibits hypothalamic specification, and likely serves to constrain the relative size of the hypothalamic region prior to the onset of neurogenesis.

## Discussion

This study provides a comprehensive roadmap for the molecular mechanisms controlling hypothalamic induction, regionalization, and initiation of neurogenesis, made possible by the use of the readily accessible and more slowly developing chick model system. It is not surprising, given the central role of the hypothalamus in the regulation of many evolutionarily ancient innate behaviors, that the overall spatial organization of the chick hypothalamus is highly similar to that of the mammalian hypothalamus during the early stages of neurogenesis. Given this finding, this study provides unprecedented insight into processes that orchestrate initial hypothalamic patterning and regionalization that are likely to be broadly evolutionarily conserved.

Owing to its complicated anatomy and rapid growth, the organization of the early developing hypothalamus, and the location of hypothalamic progenitor cells relative to other forebrain structures, has historically been controversial. New molecular markers identified here show that the HH8 hypothalamus closely resembles a floor plate-like domain sitting directly adjacent to prethalamus-like tissue. Studies in zebrafish show that the prethalamus is the earliest-developing forebrain structure (Staudt and Houart, 2007), and indeed *OLIG2*-expressing cells can be identified in the prospective prethalamus as early as HH8. Transiently (between HH8-HH10) *OLIG2*-positive cells are found concentrically surrounding the developing hypothalamic floor plate, including a subset within the high *SIX6*, *RAX*-positive region, apparently prefiguring the position of the anterior ID. Our study provides clear evidence to support the idea that the hypothalamus arises from prethalamus-like tissue.

Furthermore, our study demonstrates that FST derives from prethalamic-like neuroepithelium and actively constrains the extent of the developing hypothalamus. Mutant analysis has shown that *NKX2-1* promotes hypothalamic identity while simultaneously repressing prethalamic identity (Kim et al., 2020), and FST may mediate its effects in part by inhibiting a factor responsible for inducing *NKX2-1* expression. However, the direct mechanism of action of FST is unknown. A strong candidate for this factor is prechordal mesoderm-derived Nodal, which has been widely shown to induce hypothalamic midline cells (Hong et al., 2020; Mathieu et al., 2002; Patten et al., 2003; Placzek and Briscoe, 2005). However, our explant cultures include no prechordal mesoderm, and no Nodal expression is detected in hypothalamic cells at any time point analyzed using scRNA-Seq. This raises the possibility that another target of FST may mediate these effects. One such candidate is BMP2, which is strongly expressed in a complementary pattern to FST at HH8 and HH10 (Fig. 7F’). Previous studies have shown that BMP2, BMP4, and BMP7 interact with Shh to induce and pattern the hypothalamus (Dale et al., 1997; Manning et al., 2006; Ohyama et al., 2005), while in the posterior neural tube, FST is known to modulate the level of BMP signaling to contribute to dorsoventral neural tube patterning, in conjunction with SHH (Liem et al., 2000). The complex interactions between BMPs, SHH, and FST are likely to underlie hypothalamic induction, and further studies will be required to dissect the relative contribution of these factors, and to determine the precise mechanism of action of FST.

Previous studies have shown that the hypothalamus is induced at HH8 and exists as a floor plate-like structure (Dale et al., 1997, 1999). Here we markedly extend the characterization of these cells. The HH8 hypothalamus, defined here through *FOXA2/SIX6/DBX1/NKX2-1* expression, undergoes a dramatic expansion from this point onward and appears to give rise to the majority of the basal hypothalamus (schematized in Fig. 2I,J; Fig. 3R; Fig. 4A). By HH10, both scRNA-Seq and HCR analysis reveal the existence of two spatially and molecularly distinct clusters of hypothalamic progenitors. The first of these initially continues to express floor plate-like markers, and eventually goes on to give rise to neurons of the posterior (mammillary/supramammillary) and, possibly, the anterodorsally-located paraventricular nucleus. The second -- which expresses relatively high levels of *SIX6*, *RAX,* and *FGF10* -- gives rise to neurons of the tuberal hypothalamus, including the ventromedial and arcuate nuclei.

The possibility of a shared developmental program between the posterior (mammillary/supramammillary) and paraventricular hypothalamus is surprising, given that they are located at opposite ends of the hypothalamus. At HH20, posterior and paraventricular hypothalamic neural precursors cluster together, while HCR analysis shows that arginine-vasopressin (*AVP*; expressed by paraventricular neurons) is found in the anterior mammillary region (Fig. 6G, H) (in agreement with previous studies on later stage chicken embryos ((Caqueret et al., 2005)) as well as the paraventricular region. It has been previously reported that progenitors at the anterior and posterior borders of the mouse hypothalamus coexpress a number of genes, including *LHX5*, *SIM1,* and *NEUROG2* (Kim et al., 2020; Shimogori et al., 2010); this work further emphasizes the deep similarity of these neurons. While similar neuronal differentiation programs may mask underlying differences in progenitor origin, an intriguing possibility, raised by our studies, is that AVP-expressing neurons detected adjacent to *VAX1/RAX*-positive progenitors in the antero-dorsal hypothalamus have migrated into this position from mammillary regions, rather than being born from *VAX1/RAX*-positive progenitors(Mayer et al., 2018). Differential Wnt signaling, which acts to posteriorize the zebrafish and mouse hypothalamus, may ultimately distinguish posterior (mammillary/supramammillary) and paraventricular progenitors/precursors (Lee et al., 2006; Newman et al., 2018).

By HH18, the organization of the chick hypothalamus broadly resembles that of the E11.5 mouse, and clear counterparts of most major chick neuronal precursor populations are observed in E11-13 mice and GW 10 humans. Paraventricular and mammillary progenitor populations are clearly distinguished, and prethalamic markers are already extending into the basal hypothalamus in an anterior direction via the ID and ventrally via the TT. In the tuberal hypothalamus, selective markers of both ventromedial hypothalamus and arcuate nucleus, such as *NR5A1* and *POMC*, are both observed, although neither structure is yet spatially distinct.

Alongside the extensive conservation between the chick, mouse, and human developing hypothalamus, a few key differences stand out. First, we observe high, early, and selective expression of hypocretin in *NR5A1*-positive cells, which do not express hypocretin in mammals. Previous studies have reported the existence of medially-located hypothalamic hypocretin cells in a number of non-mammalian species, including chick, in addition to the lateral hypothalamic population present in mammals (Godden et al., 2014). Second, in the E11.5 mouse posterior hypothalamus, *EMX2* labels the mammillary region, *PITX2* labels both mammillary and supramammillary, while *FOXA1/2* labels only the supramammillary region (Shimogori et al., 2010). In chick, in contrast, *FOXA1*/*2*, *PITX2,* and weak *EMX2* expression coincide in a region taken to include mammillary and supramammillary precursors, although an *EMX2*-positive, *PITX2*-negative region anterior to this may also contribute towards the MMN. ScRNA-Seq identified chick *CALB2*-positive mammillary and *TAC1*-positive supramammillary neural precursors. Finally, we are unable to detect clear counterparts of either premammillary or lateral hypothalamic neural precursors, which likely reflects the earlier developmental age of the chick samples analyzed here, in comparison to E11.5 mouse and GW10 human

Previous studies have highlighted the relevance to human health of a detailed understanding of hypothalamic development. Congenital metabolic disorders and obesity can result from Mendelian mutations in transcription factors that control hypothalamic patterning (Blanchet et al., 2017; Holder et al., 2000), and mutations in genes identified as important for controlling hypothalamic and neurogenesis in this study may contribute to multigenic disorders that may have a hypothalamic origin, such as Type 2 diabetes, sleep disorders, and depression (Bao et al., 2008; Biran et al., 2015; Dearden and Ozanne, 2015). Understanding the molecular mechanisms controlling early hypothalamic development is also critical to efforts towards induced differentiation of specific hypothalamic neuronal subtypes from embryonic stem (ES) or induced pluripotent stem (iPS) cells (Merkle et al., 2015; Nagasaki et al., 2015; Seifinejad et al., 2019; Wang et al., 2016), as well as for directed reprogramming of hypothalamic glial cells (Kano et al., 2019; Yoo et al., 2021). Cell-based approaches such as these may ultimately hold the potential for the directed rewiring of core hypothalamic regulatory circuitry for treating a broad range of homeostatic disorders.

## Acknowledgments

We thank A. Fletcher, A. Furley, M. Towers, I. Barbaric, and T. Shimogori for helpful comments on the manuscript. We thank Transcriptomics and Deep Sequencing Core (Johns Hopkins) for sequencing of scRNA-Seq libraries. This work was supported by the Wellcome Trust (212247/Z/18/Z) to MP, NIH (R01DK108230) to SB, and the Maryland Stem Cell Research Fund (2019-MSCRFF-5124) to DWK.

## Data availability

All scRNA-Seq data are available on GEO, GSE171649.

**Table ST1:** Differentially expressed genes across clusters at HH8.

**Table ST2:** Differentially expressed genes across clusters at HH10.

**Table ST3:** Differentially expressed genes across clusters at HH8 & HH10.

**Table ST4:** Differentially expressed genes across clusters at HH13.

**Table ST5:** Differentially expressed genes across clusters at HH15/16.

**Table ST6:** Differentially expressed genes across clusters at HH13 & HH15/16.

**Table ST7:** Differentially expressed genes across clusters at HH18/19.

**Table ST8:** Differentially expressed genes across clusters at HH20/21.

**Table ST9:** Differentially expressed genes across clusters across the entire hypothalamic aggregate dataset.

**Table ST10:** Differentially expressed genes across clusters at HH18/19 & HH20/21.

**Index of Key Resources**: HCR probes and antibodies used.

## Methods

### Chick Collection

Fertilized Bovan Brown eggs (Henry Stewart & Co., Norfolk, UK) were used for all experiments. All experiments were performed according to relevant regulatory standards (University of Sheffield). Eggs were incubated and staged according to the Hamburger-Hamilton chick staging system (Hamburger and Hamilton, 1992).

### Tissue collection

Hamburger-Hamilton stage HH8, HH10, HH13/14, HH15/16, HH18/19, and HH20/21 chick embryos were harvested and neuroectoderm was isolated after Dispase I (Cat No. 4942086001, Roche) treatment (Pearson et al., 2011). Hypothalamic tissue and small amounts of neighboring tissue were manually dissected at each stage, pooled, and transferred to Hibernate E medium (Cat No. HE500, BrainBits LLC). Tissue was enzymatically digested using Papain (2 mg/ml, Cat No. LS003119, Worthington Chemicals), and cells dissociated into single cells using fire-polished glass pipettes. Papain was neutralized with Hibernate-E medium with B27 and GlutaMAX (Kim et al., 2020). Cells were pelleted at 500g at 4°C, washed twice in 1 ml cold 1 x PBS, and then resuspended in 200 ul chilled 1 x PBS by gentle pipetting. Cells were then fixed with 800 μl cold 100% methanol as previously described (Alles et al., 2017). Cells were stored at -80°C until scRNA-Seq.

### scRNA-Seq Data generation

Methanol-fixed chick embryos of dissected hypothalamus at HH8, HH8, HH13, HH15/16, HH18/19, HH20/21 were re-hydrated as previously described (Alles et al., 2017). Briefly, cells were washed twice (3000 x g for 5 min at 4°C) and resuspended in RNAse-free PBS with 1% BSA and 0.5 U/ul RNase inhibitor (Cat. N2615, Promega). Cells were then used for the 10x Genomics Chromium Single Cell System (10x Genomics, CA, USA) using V3.0 chemistry per manufacturer’s instruction, loading between 8,000 and 12,000 cells per run. Libraries were sequenced on Illumina NextSeq 500 with ∼ 200 million reads per library. Sequenced files were processed through the CellRanger pipeline (v.3.10, 10x Genomics) using the Ensembl *Gallus gallus* genome (GRCg6a, release 95) compiled with the CellRanger *mkref* function.

### scRNA-Seq Data analysis

Seurat v3 (Butler et al., 2018) was used to perform analysis following the pipeline described previously (Kim et al., 2020), selecting cells with more than 1000 genes and 1000 UMI, normalized using Seurat *scTransform*, and Harmony (Korsunsky et al., 2019) was used to regress batch effects. Louvain algorithm with a default parameter was used to generate scRNA-Seq clusters, and HCR images across chicken developmental stages and previous hypothalamic scRNA-Seq database HyDD (Kim et al., 2020) were used to identify the spatial location and identity of all hypothalamic-derived clusters in each developmental stage. Extra-hypothalamic clusters were identified when the key gene markers of the cluster were not expressed in HyDD and validated using HCR and by consulting GEISHA (Antin et al., 2014). Differential gene expression tests were performed using Seurat *FindAllMarkers* (LR, regressing depth and cell number variance).

RNA velocity (La Manno et al., 2018) was used to understand the potential developmental trajectory of chicken hypothalamus development and to verify 1) migration of presumptive prethalamus-like cells into the hypothalamus, 2) developmental trajectories of mammillary bodies, PVH/SON, and tuberal hypothalamus. Kallisto and Bustools (Melsted et al., 2021) were used to obtain spliced and unspliced transcripts using --lamanno with GRCg6a chicken genome. Scanpy (Wolf et al., 2018) and scVelo (Bergen et al., 2020) were used to process the Kallisto output with the following parameters (parameters), based on UMAP coordinates obtained from Seurat.

Monocle v3 (Trapnell et al., 2014) was used to perform pseudotime analysis (q value < 0.001) to identify differences in gene expression across prethalamus, tuberal, mammillary, and PVN/SON development identified from RNA velocity analysis.

NicheNet (Browaeys et al., 2020) was used to identify receptor-ligand interaction between HH8 and HH10, and HH13 and HH15, accounting that any cluster can be receiver and sender. First, key genes expressed in the datasets (more than 10% of the population) and background genes were identified. NicheNet ligand activity analysis was then performed by ranking putative ligands and receptors were then inferred from the selected gene lists, and ligand-receptor interaction scores were calculated based on validated ligand-receptor database.

To identify the correlation between chicken, mouse, and human, post-mitotic hypothalamic cells from HH18 and HH20 (this work), E11-E13 mouse scRNA-Seq (Kim et al., 2020), and gestational week 10 human scRNA-Seq (Zhou et al., 2020) were used. Annotation of the human scRNA-Seq dataset was based on a mouse scRNA-Seq database and as shown in the original publication. Key markers defining individual spatially identified hypothalamic nuclei of the chicken, mouse, and human were then used to plot correlations with genesorteR (Ibrahim and Kramann).

### Explant culture

Explants of prospective hypothalamus were isolated from either HH4 or HH6 embryos by Dispase treatment and cultured in collagen beds (Ohyama et al., 2005). Explants were either treated with recombinant FST (7.5 μg/ml, Cat no. 769-FS-025, R&D Systems) or anti-FST antibody (20 μ/ml, AF669, R&D Systems) for 42 hr, and processed for *in situ* hybridization chain reaction (HCR).

### *In vivo* manipulation of follistatin signaling

HH5-6 embryos were injected with 5 ul recombinant FST (7.5 ug/ml) or anti-FST antibody (20 ug/ml) and allowed to develop to HH14 (42 hrs), and processed for HCR.

### Immunohistochemistry

Explants were analyzed by immunohistochemistry according to standard techniques (Manning et al., 2006). Following cryosectioning, the sections were analyzed with the following antibodies anti-NKX2.1 (1:2000, (Ohyama et al., 2005)) and anti-Pax6 (1:50, DSHB). Secondary antibodies (1:500, Jackson Immunoresearch) were conjugated with Cy3 or FITC. Images were taken using a Zeiss Apotome or Olympus BX60 and Spot RT software v3.2.

### Chicken HCR

Hamburger & Hamilton stage 8-20 embryos were harvested and fixed in 4% Paraformaldehyde. HCR v3.0 was performed on embryos and cryosections using reagents and modified protocol from Molecular Instruments, Inc.. Samples were preincubated with a hybridization buffer for 30 min and the probe pairs were added and incubated at 37°C overnight. The next day samples were washed 4 times in the probe wash buffer and 2 times in the 5x SSC buffer and preincubated in Amplification buffer for 5 min. Even and odd hairpins for each of the genes were snap-cooled by heating at 95°C for 90 sec and cooling to RT for 30 min. The hairpins were then added to the amplification buffer and added to the samples and incubated overnight at RT in dark. Samples were then washed in 5x SSC and DAPI was added as a counterstaining. For multiplexing, after imaging with the first set of probes, the slides were treated with DNAase (0.05 U/µl), washed 3 times in 30% Formamide and 2x SSC and 3 times in 2x SSC. Slides were then preincubated with the hybridization buffer, the next set of probes were added, and the process was repeated.

---

**Figure S1.**
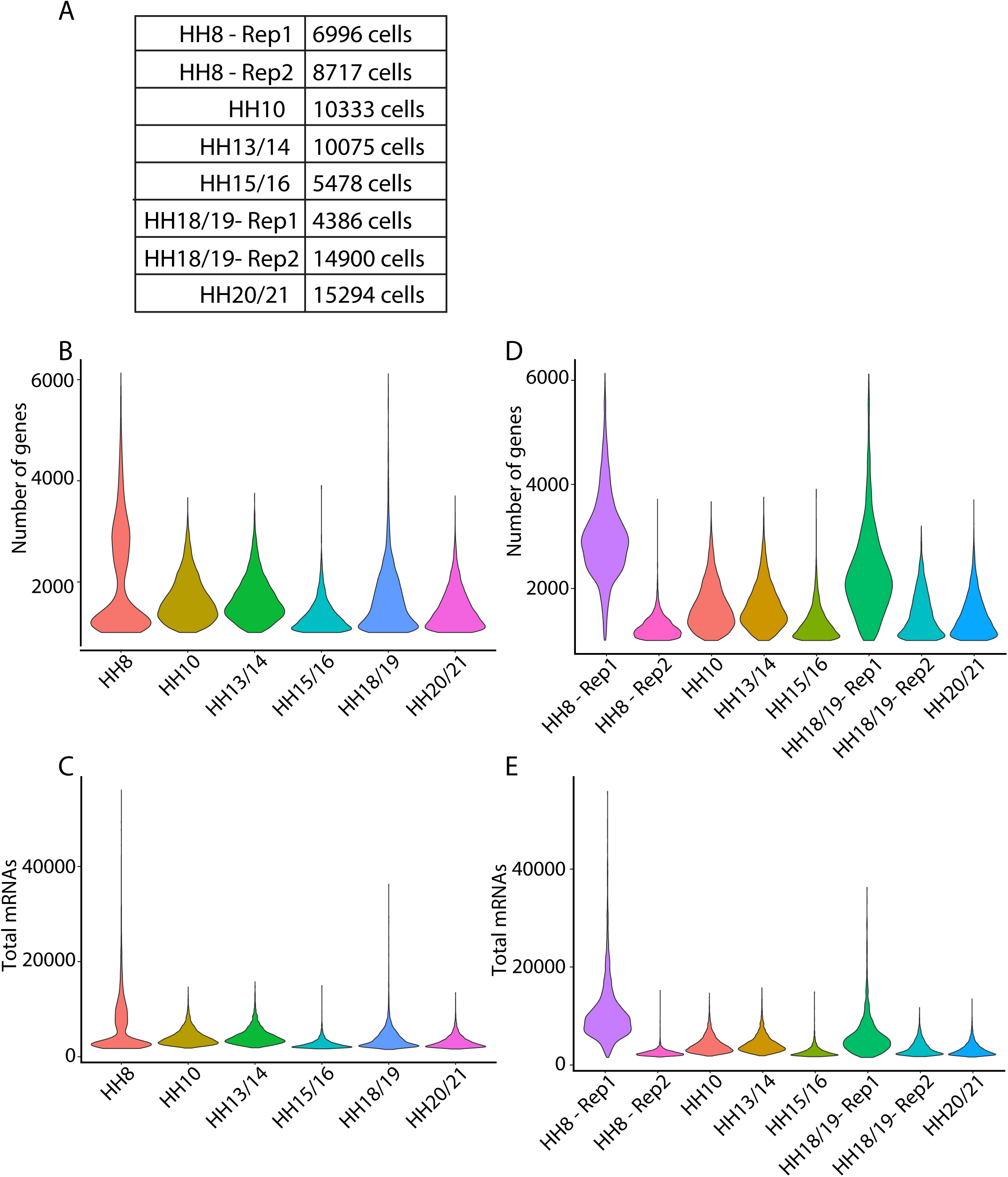
(A) Number of cells per scRNA-Seq library, (B-E) Mean number of genes (B, D), and mean number of total mRNAs (UMI) (C, E) across the entire chicken time-points (B, C), and across scRNA-Seq libraries (D, E).

**Figure S2.**
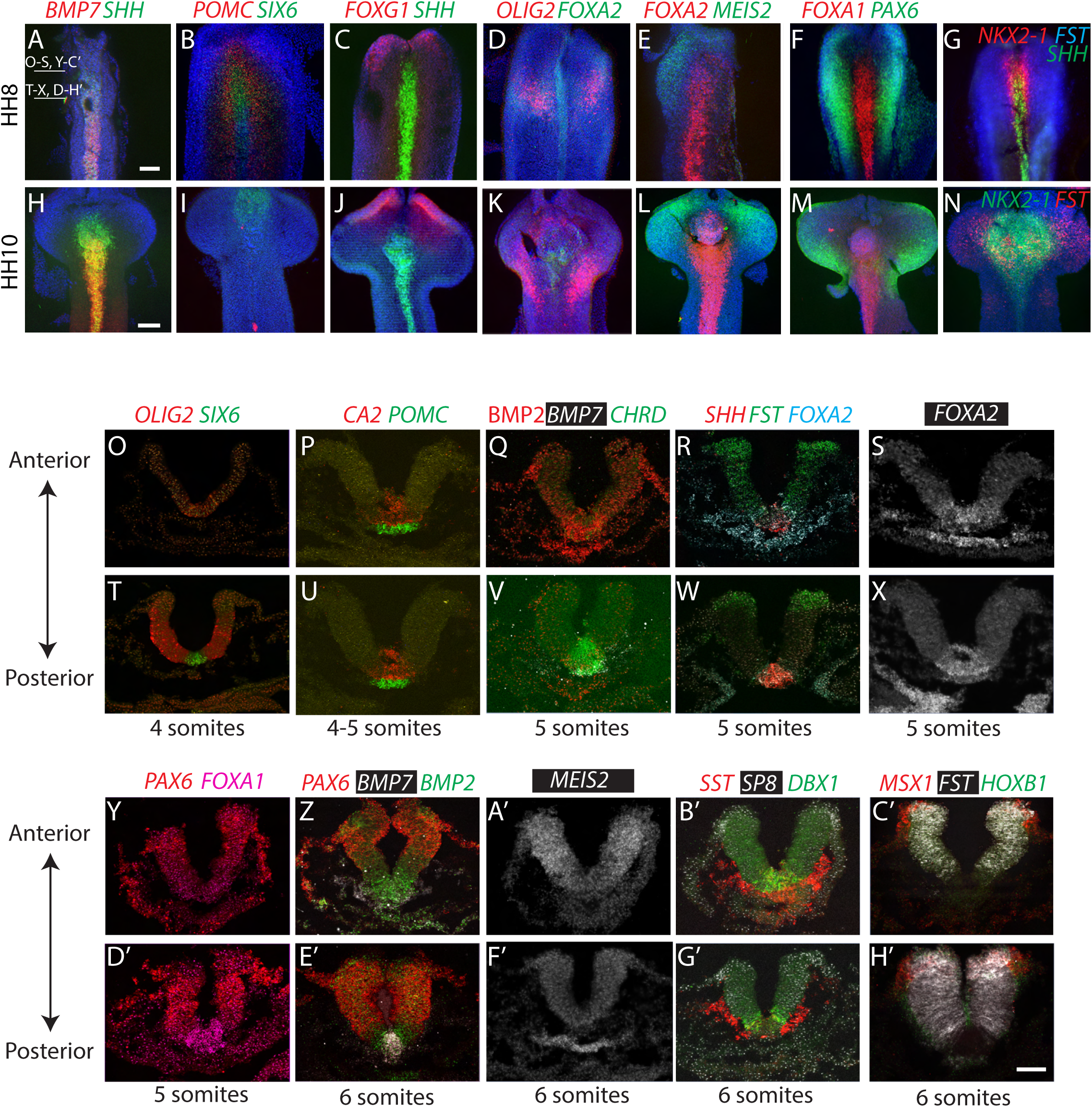
(A-N) Maximum intensity projections after wholemount HCR *in situ* hybridization on isolated neuroectoderm at HH8 (A-G) and HH10 (H-N) showing *BMP7*/*SHH* (A, H), *POMC/SIX6* (B, I), *FOXG1/SHH* (C, J), *OLIG2/FOXA2* (D, K), *FOXA2/MEIS2* (E, L), *FOXA1/PAX6* (F, M), *NKX2-1/FST*/*SHH* (G) or *NKX2-1/FST* (N). (O-H’) HCR *in situ* hybridization of transverse sections at HH8, 200 μm (O-S, Y-C’) or 350 μm (T-X, D’-H’) posterior to the neuropore showing *OLIG2*/*SIX6* (O, T), *CA2*/*POMC* (P, U), *BMP2*/*BMP7*/*CHRD* (Q, V), *SHH*/*FST*/*FOXA2* (R, W), *FOXA2* (S, X), *PAX6*/*FOXA1* (Y, D’), *PAX6*/*BMP7*/*BMP2* (Z, E’), *MEIS2* (A’, F’), *SST*/*SP8/DBX1* (B, G’), *MSX1*/*FST*/*HOXB1* (C’, H’).

**Figure S3.**
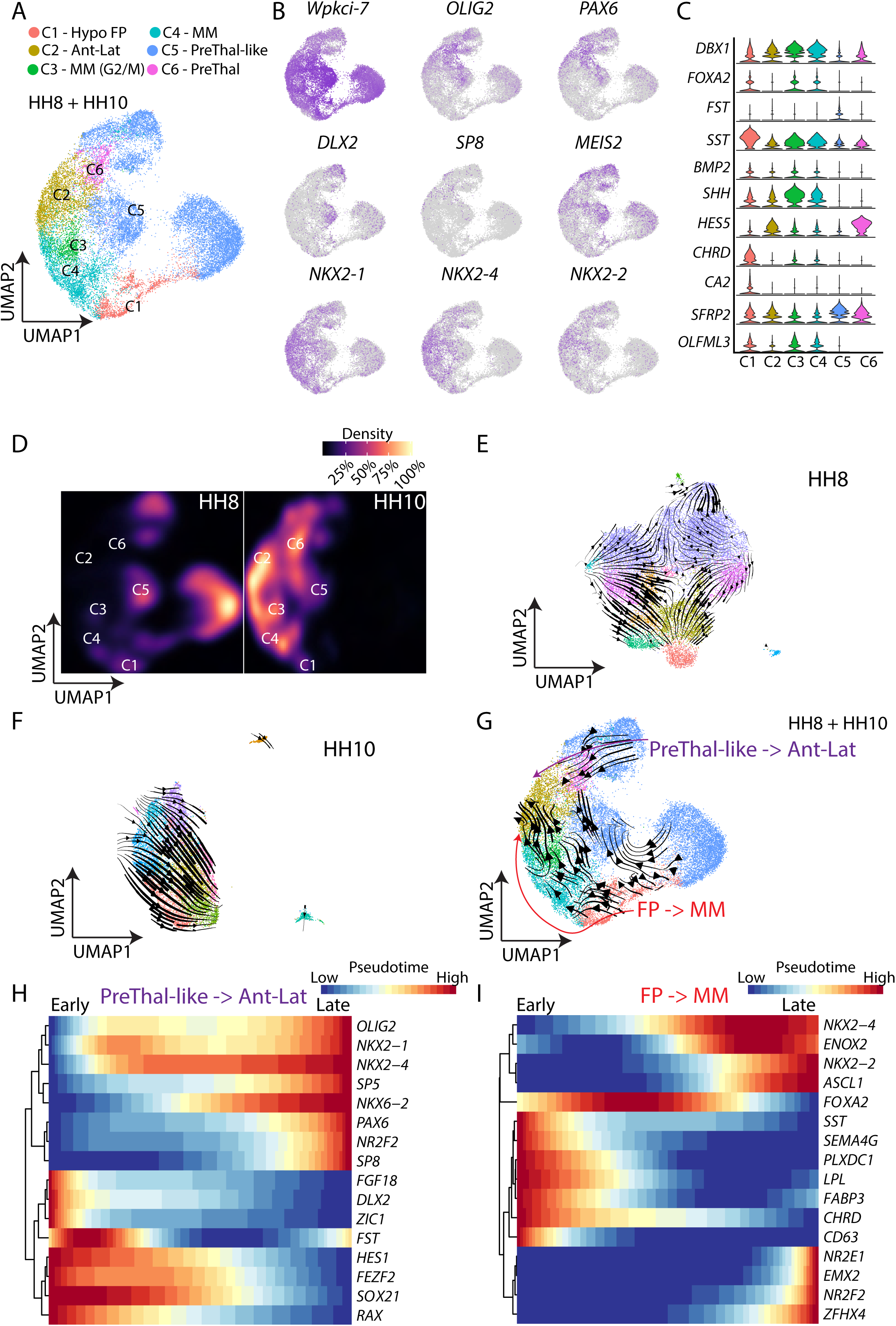
(A) UMAP plot showing combined HH8/HH10 datasets. (B) UMAP plot showing expression of prethalamic and hypothalamic (floor plate-like) genes. (C) Violin plot showing expression of key genes of HH8/HH10 datasets. (D) Heatmap plot showing the distribution of HH8 and HH10 cells from figure panel A. (E) UMAP plot showing HH8 scRNA-Seq with RNA velocity trajectories. (F) UMAP plot showing HH10 scRNA-Seq with RNA velocity trajectories. (G) UMAP plot showing HH8/HH10 scRNA-Seq trajectories obtained from RNA velocity. (H-I) Pseudotime analysis showing changes in gene expression, indicating conversion of prethalamus to anterior/tuberal/lateral hypothalamus (H), and floor-plate into posterior hypothalamus (I). Ant = Anterior, FP = Floor plate, Hypo = Hypothalamus, Lat = Lateral, MM = Mammillary, PreThal-like = Prethalamus-like progenitors, PreThal = Prethalamus.

**Figure S4.**
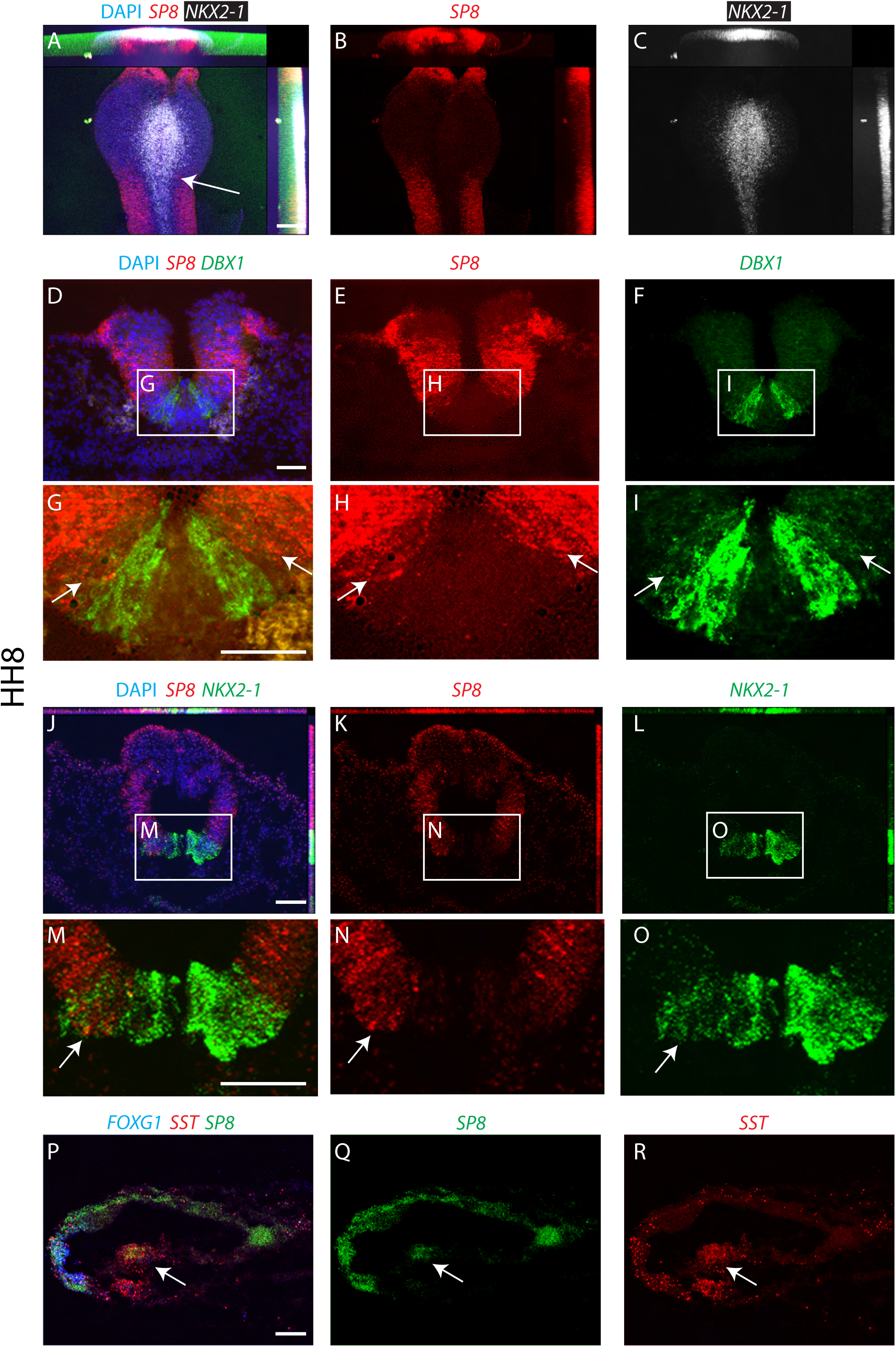
HCR in situs showing co-expression of hypothalamic and prethalamic-like markers. (A-C) Wholemount view of isolated neuroectoderm at HH9 showing *SP8* (A,B) and *NKX2-1* (A,C). (D-I) Transverse sections at HH8 showing *SP8* (D,E,G,H) and *DBX1* (D,F,G,I). (J-O) Transverse sections at HH8 showing *SP8* (J,K,M,N) and *NKX2-1* (J,L,M,O). (P-R) Parasagittal section at HH8 showing *SP8* (P,Q) and *SST* (P,R). Arrow in (A) indicates approximate position of section planes in (D-R). Arrows in (G-I,M-R) point to co-expressing cells. Scale bar = 100 μm.

**Figure S5.**
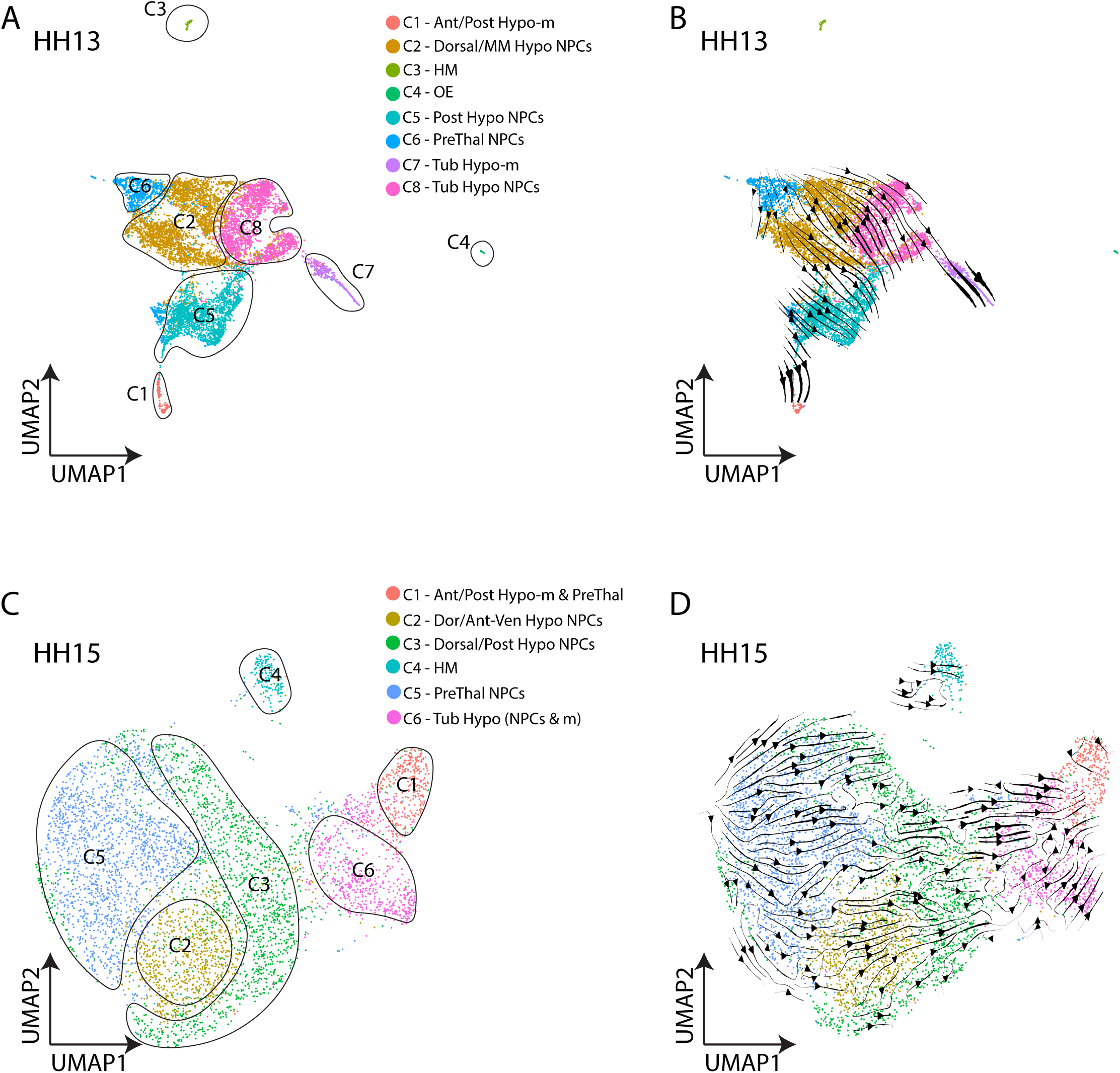
(A-B) UMAP plot showing HH13 scRNA-Seq clusters (A) and RNA velocity trajectories (B). (C-D) UMAP plot showing HH15 scRNA-Seq clusters (C) and RNA velocity trajectories (D). Ant = Anterior, Hypo = Hypothalamus, Hypo-m = Maturing hypothalamus, HM = Head mesoderm, MM = Mammillary, NPCs = Neural precursor cells, OE = Oral ectoderm, Post = Posterior, PreThal = Prethalamus, Tub = Tuberal, Ven = Ventral

**Figure S6.**
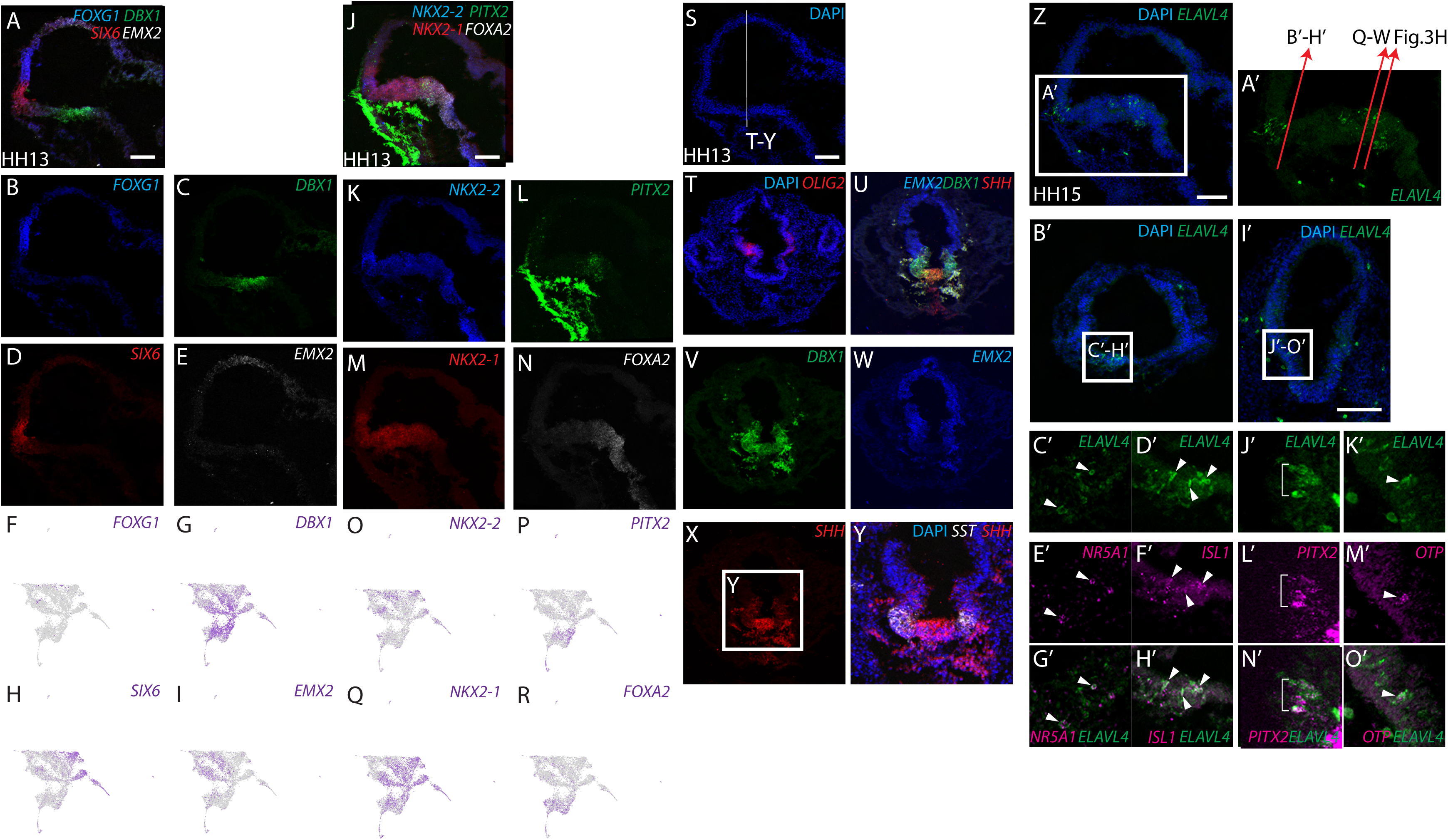
(A-E) HCR *in situ* hybridization at HH13 (parasagittal plane) showing *FOXG1* (A, B), *DBX1* (A, C), *SIX6* (A, D), *EMX2* (A, E). UMAP plot showing gene expression of *FOXG1* (F), *DBX1* (G), *SIX6* (H), *EMX2* (I). (J-N) HCR *in situ* hybridization at HH13 (parasagittal plane) showing *NKX2-2* (J, K), *PITX2* (J, L), *NKX2-1* (J, M), *FOXA2* (J, N). UMAP plot showing gene expression of *NKX2-2* (O), *PITX2* (P), *NKX2-1* (Q), *FOXA2* (R). (S-Y) HCR *in situ* hybridization at coronal planes (T-Y) at HH13 (S, parasagittal plane) showing *OLIG2* (T), *EMX2* (U, W), *DBX1* (U, V), *SHH* (U, X, Y), and *SST* (Y). HCR *in situ* hybridization at coronal planes (B’-O’) at HH15 (Z’-A’, parasagittal plane) showing *ELAVL4* (B’-D’, G’-K’, N’, O’), *NR5A1* (E’, G’), *ISL1* (F’, H’), *PITX2* (L’, N’), *OTP* (M’, O’). Scale bar = 100 μm.

**Figure S7.**
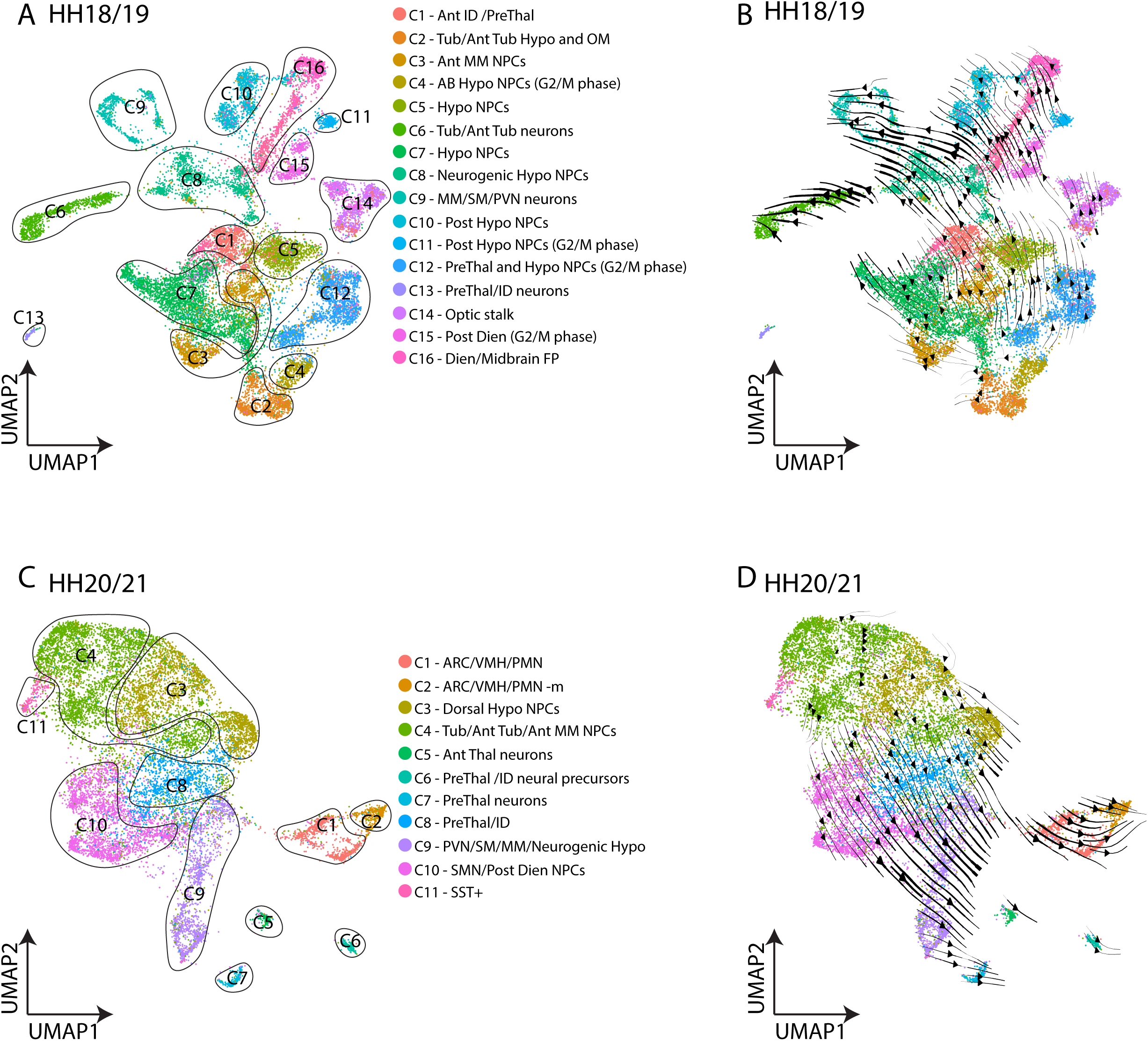
(A-B) UMAP plot showing HH18/19 scRNA-Seq clusters (A) and RNA velocity trajectories (B). (C-D) UMAP plot showing HH20/21 scRNA-Seq clusters (C) and RNA velocity trajectories (D). ARC = Arcuate nucleus of the hypothalamus, Ant = Anterior, Dien = Diencephalon, FP = Floor plate, Hypo = Hypothalamus, ID = Intrahypothalamic diagonal, MM = Mammillary, NPC = Neural precursor cells, OM = Optic midline, PreThal = Prethalamus, PMN = Premammillary nucleus, Post = Posterior, PVN = Paraventricular nucleus of the hypothalamus, SM = Supramammillary, Tub = Tuberal, VMH = Ventromedial hypothalamus.

**Figure S8.**
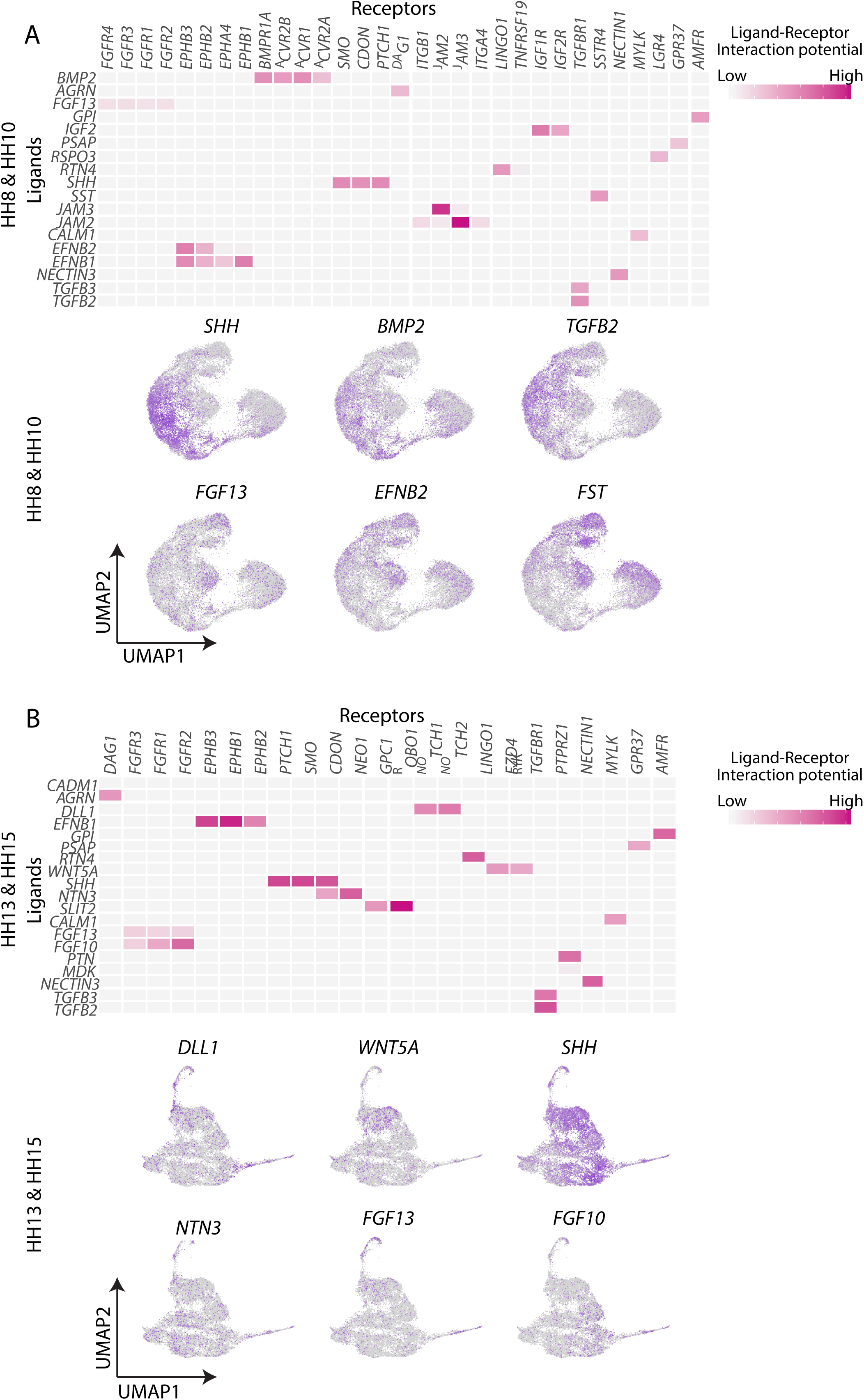
(A) Heatmap showing the interaction of ligands and known-receptors at HH8/HH10 (top), and UMAP plot showing expression of ligands at HH8/HH10 (bottom). (B) Heatmap showing the interaction of ligands and known-receptors at HH13/HH15 (top), and UMAP plot showing expression of ligands at HH13/HH15 (bottom).

## KEY RESOURCES TABLE

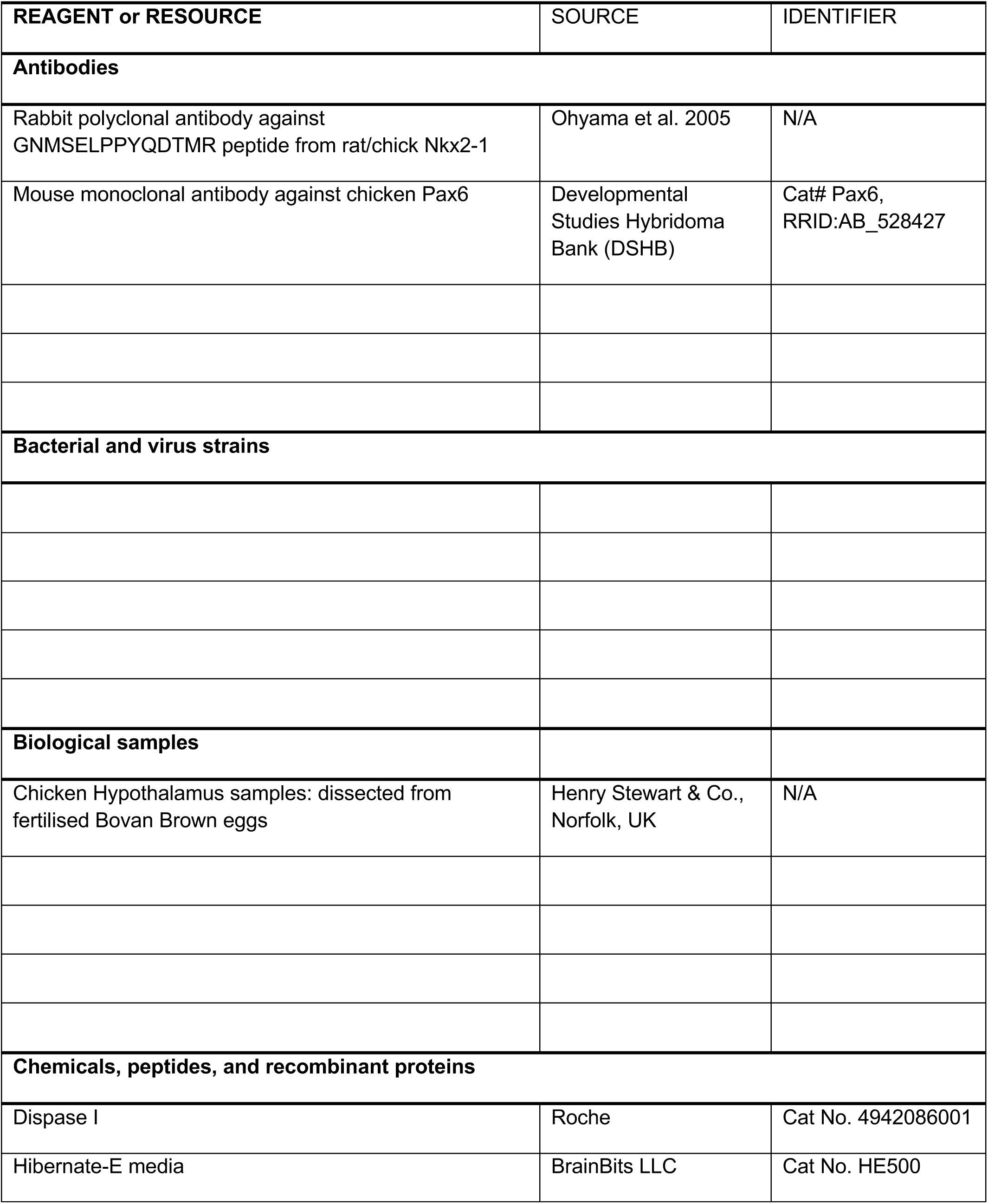
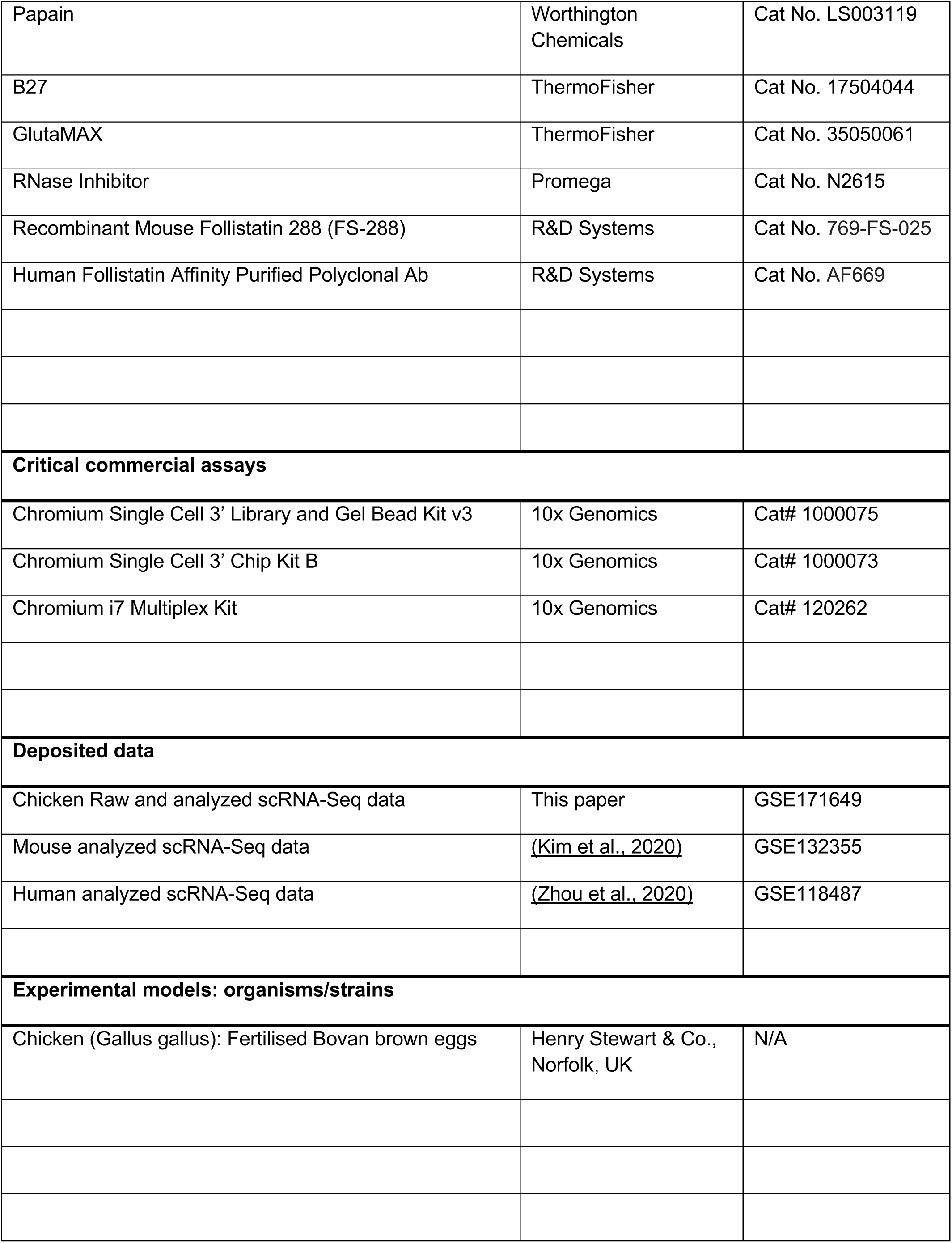
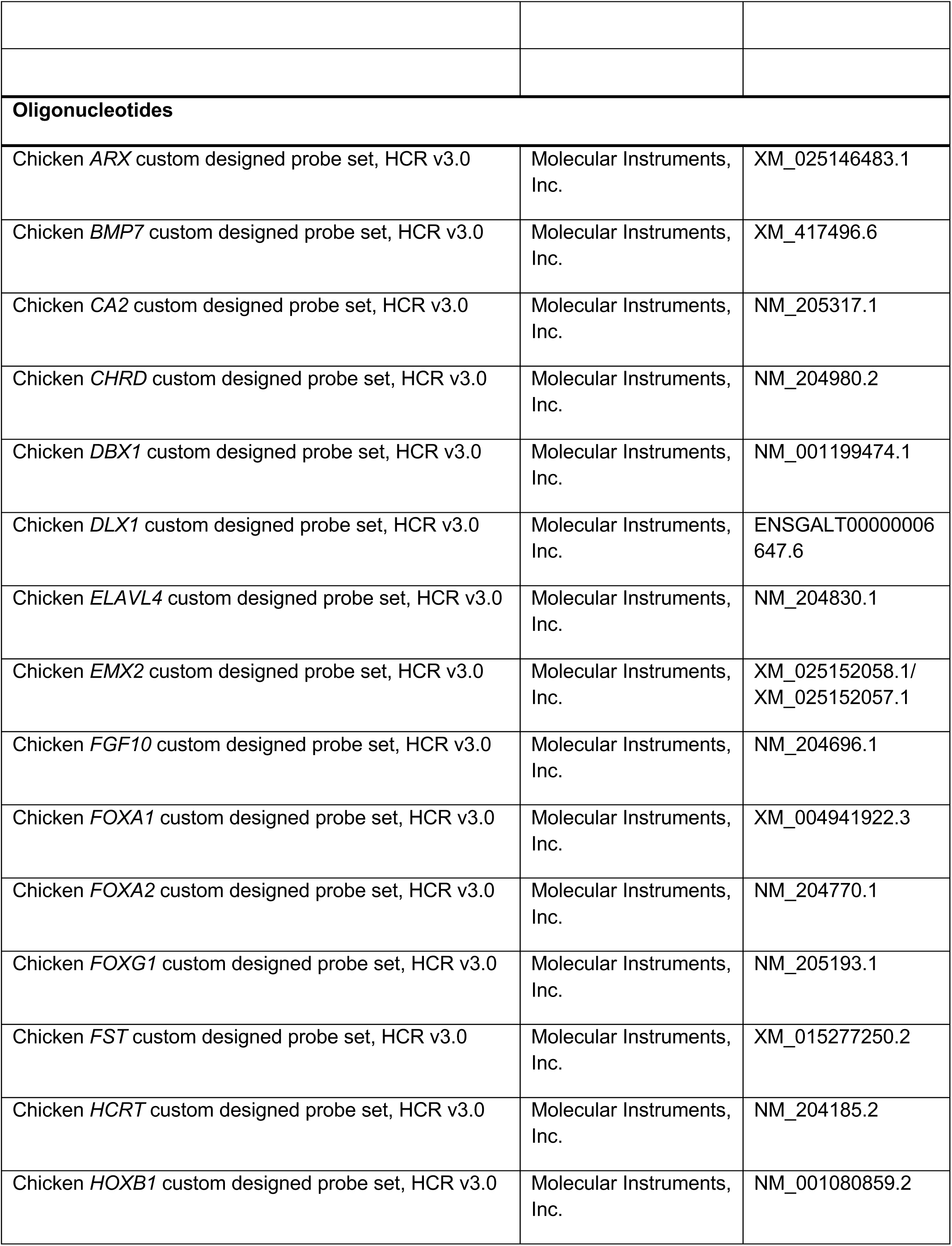
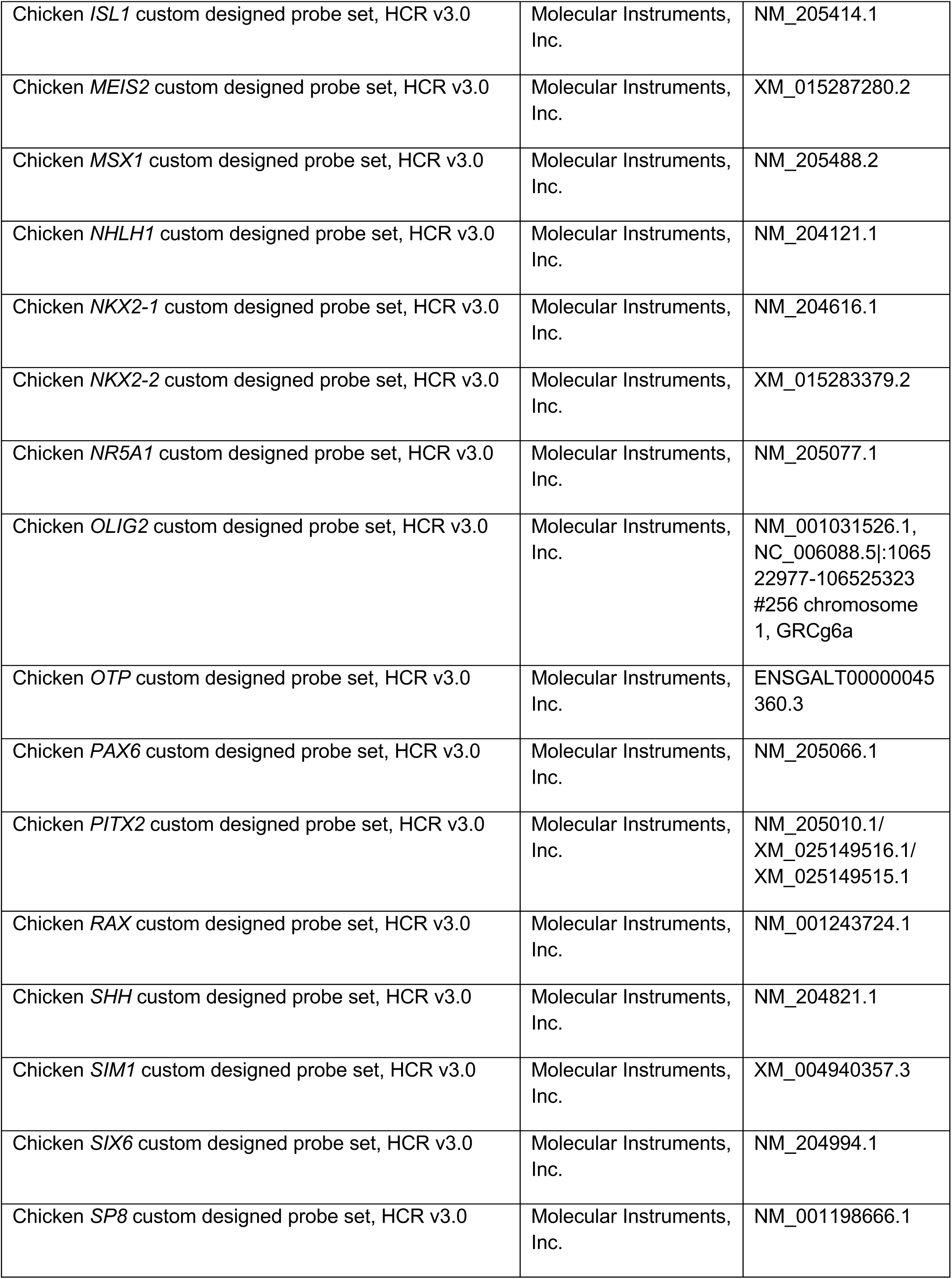
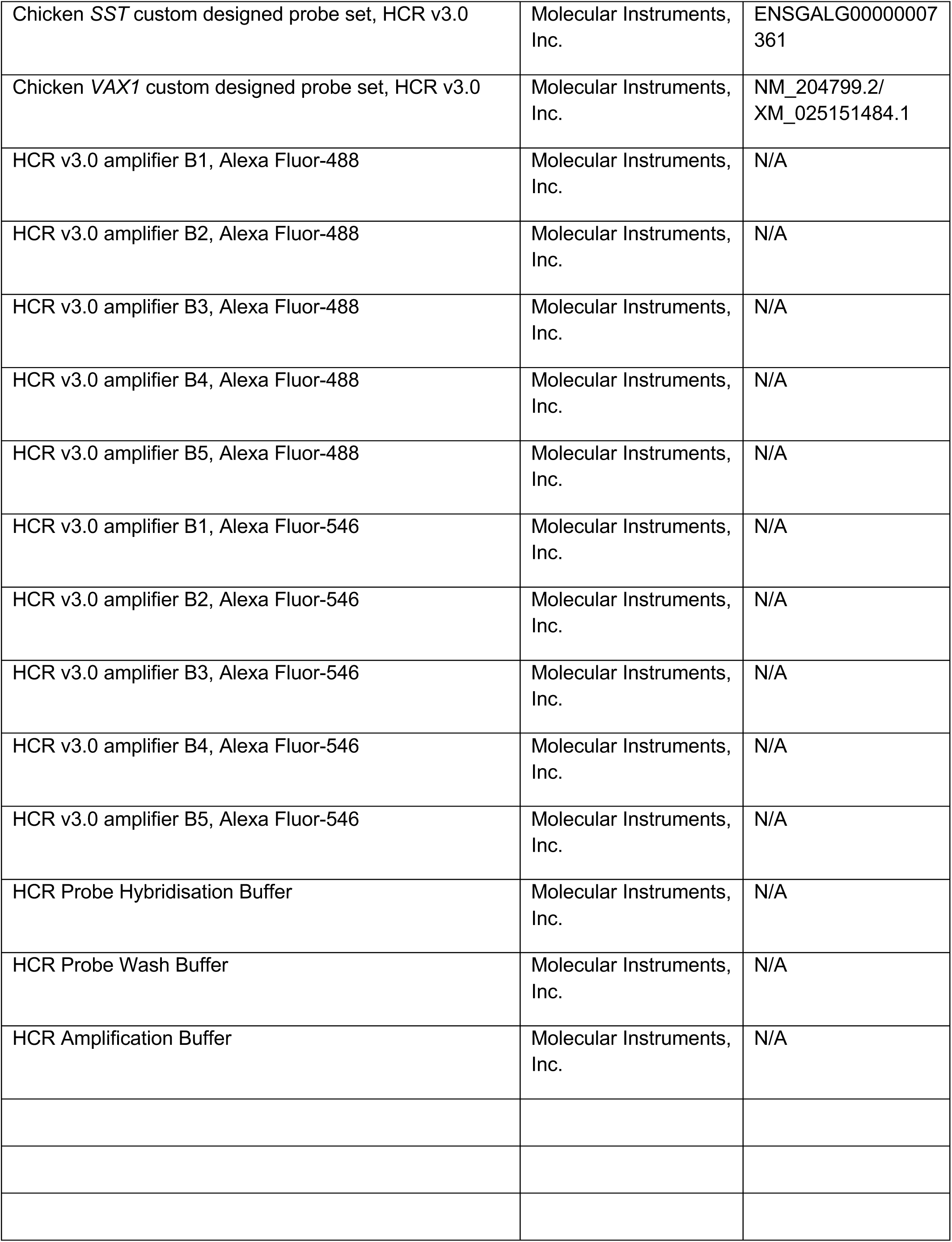
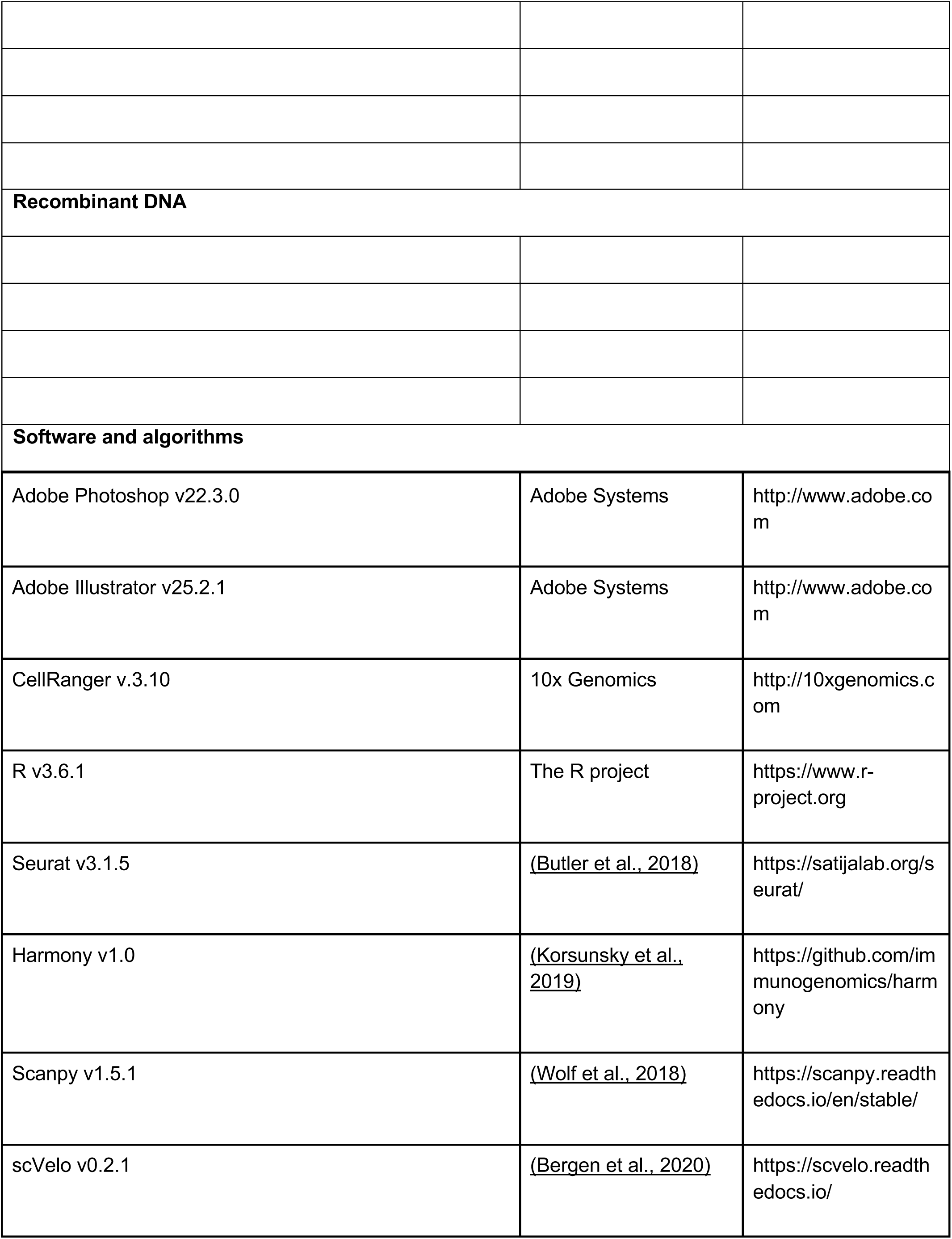
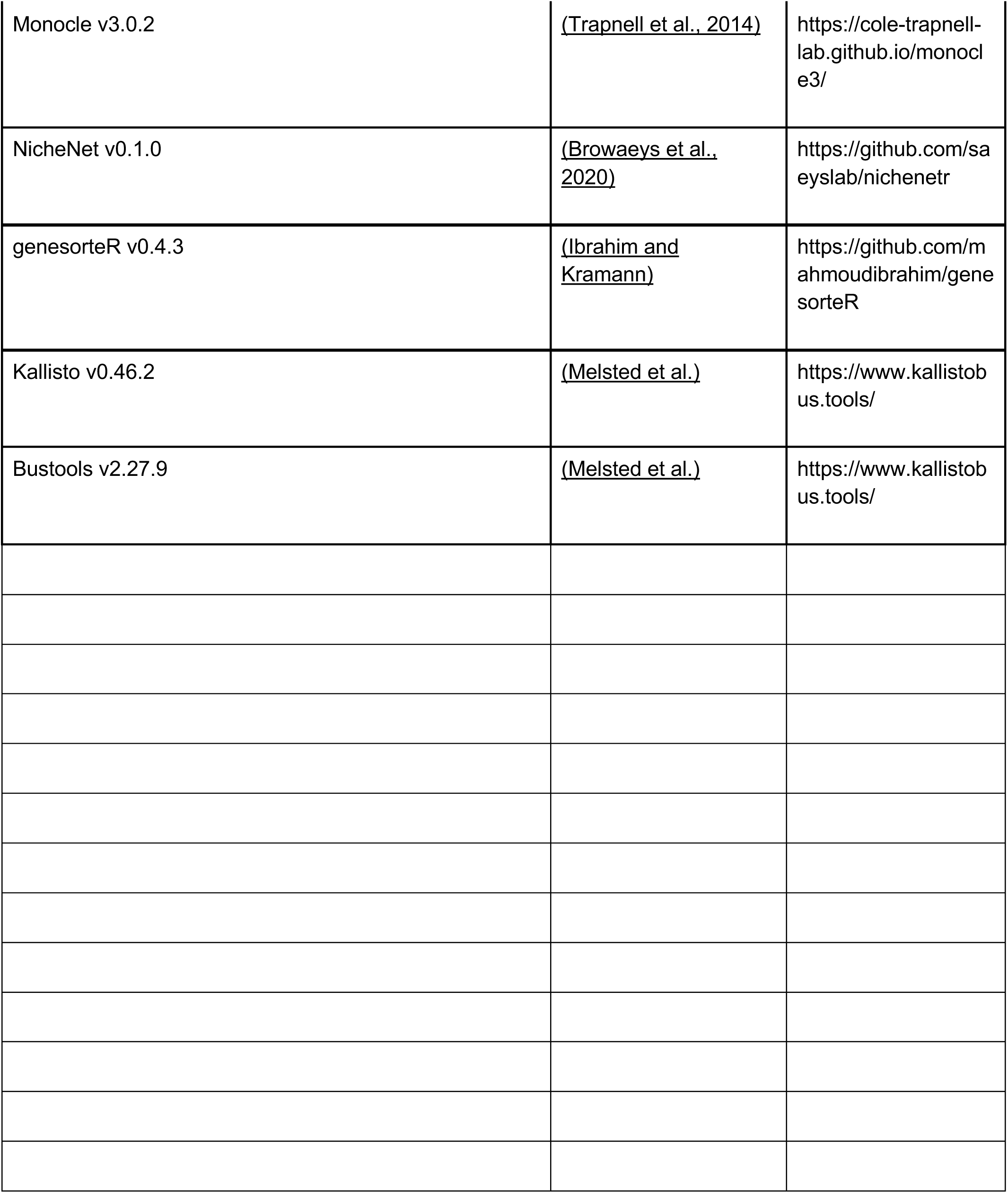

## References

1. Alles, J., Karaiskos, N., Praktiknjo, S.D., Grosswendt, S., Wahle, P., Ruffault, P.-L., Ayoub, S., Schreyer, L., Boltengagen, A., Birchmeier, C., et al. (2017). Cell fixation and preservation for droplet-based single-cell transcriptomics. BMC Biol. 15, 44.

2. Antin, P.B., Yatskievych, T.A., Davey, S., and Darnell, D.K. (2014). GEISHA: an evolving gene expression resource for the chicken embryo. Nucleic Acids Res. 42, D933–D937.

3. Aslanpour, S., Han, S., Schuurmans, C., and Kurrasch, D.M. (2020). Neurog2 acts as a Classical Proneural Gene in the Ventromedial Hypothalamus and Is Required for the Early Phase of Neurogenesis. J. Neurosci. 40, 3549–3563.

4. Aujla, P.K., Naratadam, G.T., Xu, L., and Raetzman, L.T. (2013). Notch/Rbpjκ signaling regulates progenitor maintenance and differentiation of hypothalamic arcuate neurons. Development 140, 3511–3521.

5. Bao, A.-M., Meynen, G., and Swaab, D.F. (2008). The stress system in depression and neurodegeneration: focus on the human hypothalamus. Brain Res. Rev. 57, 531–553.

6. Bedont, J.L., LeGates, T.A., Slat, E.A., Byerly, M.S., Wang, H., Hu, J., Rupp, A.C., Qian, J., Wong, G.W., Herzog, E.D., et al. (2014). Lhx1 controls terminal differentiation and circadian function of the suprachiasmatic nucleus. Cell Rep. 7, 609–622.

7. Bedont, J.L., Newman, E.A., and Blackshaw, S. (2015). Patterning, specification, and differentiation in the developing hypothalamus. Wiley Interdiscip. Rev. Dev. Biol. 4, 445– 468.

8. Bergen, V., Lange, M., Peidli, S., Wolf, F.A., and Theis, F.J. (2020). Generalizing RNA velocity to transient cell states through dynamical modeling. Nat. Biotechnol. 38, 1408– 1414.

9. Biran, J., Tahor, M., Wircer, E., and Levkowitz, G. (2015). Role of developmental factors in hypothalamic function. Frontiers in Neuroanatomy 9.

10. Blackshaw, S., Scholpp, S., Placzek, M., Ingraham, H., Simerly, R., and Shimogori, T. (2010). Molecular pathways controlling development of thalamus and hypothalamus: from neural specification to circuit formation. J. Neurosci. 30, 14925–14930.

11. Blanchet, P., Bebin, M., Bruet, S., Cooper, G.M., Thompson, M.L., Duban-Bedu, B., Gerard, B., Piton, A., Suckno, S., Deshpande, C., et al. (2017). MYT1L mutations cause intellectual disability and variable obesity by dysregulating gene expression and development of the neuroendocrine hypothalamus. PLoS Genet. 13, e1006957.

12. Brodie-Kommit, J., Clark, B.S., Shi, Q., Shiau, F., Kim, D.W., Langel, J., Sheely, C., Schmidt, T., Badea, T., Glaser, T., et al. (2021). Atoh7-independent specification of retinal ganglion cell identity. Science Advances 7, eabe4983.

13. Browaeys, R., Saelens, W., and Saeys, Y. (2020). NicheNet: modeling intercellular communication by linking ligands to target genes. Nat. Methods 17, 159–162.

14. Burbridge, S., Stewart, I., and Placzek, M. (2016). Development of the Neuroendocrine Hypothalamus. Compr. Physiol. 6, 623–643.

15. Butler, A., Hoffman, P., Smibert, P., Papalexi, E., and Satija, R. (2018). Integrating single-cell transcriptomic data across different conditions, technologies, and species. Nat. Biotechnol. 36, 411.

16. Caqueret, A., Coumailleau, P., and Michaud, J.L. (2005). Regionalization of the anterior hypothalamus in the chick embryo. Dev. Dyn. 233, 652–658.

17. Carter, R.A., Bihannic, L., Rosencrance, C., Hadley, J.L., Tong, Y., Phoenix, T.N., Natarajan, S., Easton, J., Northcott, P.A., and Gawad, C. (2018). A Single-Cell Transcriptional Atlas of the Developing Murine Cerebellum. Curr. Biol. 28, 2910–2920.e2.

18. Chen, X., Wyler, S.C., Li, L., Arnold, A.G., Wan, R., Jia, L., Landy, M.A., Lai, H.C., Xu, P., and Liu, C. (2020). Comparative Transcriptomic Analyses of Developing Melanocortin Neurons Reveal New Regulators for the Anorexigenic Neuron Identity. J. Neurosci. 40, 3165–3177.

19. Clark, B.S., Stein-O’Brien, G.L., Shiau, F., Cannon, G.H., Davis-Marcisak, E., Sherman, T., Santiago, C.P., Hoang, T.V., Rajaii, F., James-Esposito, R.E., et al. (2019). Single-Cell RNA-Seq Analysis of Retinal Development Identifies NFI Factors as Regulating Mitotic Exit and Late-Born Cell Specification. Neuron 102, 1111–1126.e5.

20. Cobos, I., Shimamura, K., Rubenstein, J.L., Martínez, S., and Puelles, L. (2001). Fate map of the avian anterior forebrain at the four-somite stage, based on the analysis of quail-chick chimeras. Dev. Biol. 239, 46–67.

21. Corman, T.S., Bergendahl, S.E., and Epstein, D.J. (2018). Distinct temporal requirements for Sonic hedgehog signaling in development of the tuberal hypothalamus. Development 145, dev167379.

22. Dalal, J., Roh, J.H., Maloney, S.E., Akuffo, A., Shah, S., Yuan, H., Wamsley, B., Jones, W.B., de Guzman Strong, C., Gray, P.A., et al. (2013). Translational profiling of hypocretin neurons identifies candidate molecules for sleep regulation. Genes Dev. 27, 565–578.

23. Dale, J.K., Vesque, C., Lints, T.J., Sampath, T.K., Furley, A., Dodd, J., and Placzek, M. (1997). Cooperation of BMP7 and SHH in the induction of forebrain ventral midline cells by prechordal mesoderm. Cell 90, 257–269.

24. Dale, K., Sattar, N., Heemskerk, J., Clarke, J.D., Placzek, M., and Dodd, J. (1999). Differential patterning of ventral midline cells by axial mesoderm is regulated by BMP7 and chordin. Development 126, 397–408.

25. Dearden, L., and Ozanne, S.E. (2015). Early life origins of metabolic disease: Developmental programming of hypothalamic pathways controlling energy homeostasis. Frontiers in Neuroendocrinology 39, 3–16.

26. Ferran, J.L., Sánchez-Arrones, L., Bardet, S.M., Sandoval, J.E., Martínez-de-la-Torre, M., and Puelles, L. (2008). Early pretectal gene expression pattern shows a conserved anteroposterior tripartition in mouse and chicken. Brain Res. Bull. 75, 295–298.

27. Fu, T., Towers, M., and Placzek, M.A. (2017). progenitors give rise to the chick hypothalamus by rostral and caudal growth and differentiation. Development 144, 3278–3288.

28. Fu, T., Pearson, C., Towers, M., and Placzek, M. (2019). Development of the basal hypothalamus through anisotropic growth. J. Neuroendocrinol. 31, e12727.

29. Godden, K.E., Landry, J.P., Slepneva, N., Migues, P.V., and Pompeiano, M. (2014). Early expression of hypocretin/orexin in the chick embryo brain. PLoS One 9, e106977.

30. Hamburger, V., and Hamilton, H.L. (1992). A series of normal stages in the development of the chick embryo. 1951. Dev. Dyn. 195, 231–272.

31. Holder, J.L., Jr, Butte, N.F., and Zinn, A.R. (2000). Profound obesity associated with a balanced translocation that disrupts the SIM1 gene. Hum. Mol. Genet. 9, 101–108.

32. Hong, M., Christ, A., Christa, A., Willnow, T.E., and Krauss, R.S. (2020). mutation and fetal alcohol converge on Nodal signaling in a mouse model of holoprosencephaly. Elife 9.

33. Ibrahim, M.M., and Kramann, R. genesorteR: Feature Ranking in Clustered Single Cell Data.

34. Kano, M., Suga, H., Ishihara, T., Sakakibara, M., Soen, M., Yamada, T., Ozaki, H., Mitsumoto, K., Kasai, T., Sugiyama, M., et al. (2019). Tanycyte-Like Cells Derived From Mouse Embryonic Stem Culture Show Hypothalamic Neural Stem/Progenitor Cell Functions. Endocrinology 160, 1701–1718.

35. Kapsimali, M., Caneparo, L., Houart, C., and Wilson, S.W. (2004). Inhibition of Wnt/Axin/beta-catenin pathway activity promotes ventral CNS midline tissue to adopt hypothalamic rather than floorplate identity. Development 131, 5923–5933.

36. Kim, D.W., Washington, P.W., Wang, Z.Q., Lin, S.H., Sun, C., Ismail, B.T., Wang, H., Jiang, L., and Blackshaw, S. (2020). The cellular and molecular landscape of hypothalamic patterning and differentiation from embryonic to late postnatal development. Nat. Commun. 11, 4360.

37. Kim, D.W., Liu, K., Wang, Z.Q., Zhang, Y.S., Bathini, A., Brown, M.P., Lin, S.H., Washington, P.W., Sun, C., Lindtner, S., et al. (2021). Gene regulatory networks controlling differentiation, survival, and diversification of hypothalamic Lhx6-expressing GABAergic neurons. Commun Biol 4, 95.

38. Korsunsky, I., Millard, N., Fan, J., Slowikowski, K., Zhang, F., Wei, K., Baglaenko, Y., Brenner, M., Loh, P.-R., and Raychaudhuri, S. (2019). Fast, sensitive and accurate integration of single-cell data with Harmony. Nat. Methods 16, 1289–1296.

39. La Manno, G., Soldatov, R., Zeisel, A., Braun, E., Hochgerner, H., Petukhov, V., Lidschreiber, K., Kastriti, M.E., Lönnerberg, P., Furlan, A., et al. (2018). RNA velocity of single cells. Nature 560, 494–498.

40. Larsen, C.W., Zeltser, L.M., and Lumsden, A. (2001). Boundary formation and compartition in the avian diencephalon. J. Neurosci. 21, 4699–4711.

41. Lee, B., Kim, J., An, T., Kim, S., Patel, E.M., Raber, J., Lee, S.-K., Lee, S., and Lee, J.W. (2018). Dlx1/2 and Otp coordinate the production of hypothalamic GHRH- and AgRP-neurons. Nat. Commun. 9, 2026.

42. Lee, J.E., Wu, S.-F., Goering, L.M., and Dorsky, R.I. (2006). Canonical Wnt signaling through Lef1 is required for hypothalamic neurogenesis. Development 133, 4451–4461.

43. Liem, K.F., Jr, Jessell, T.M., and Briscoe, J. (2000). Regulation of the neural patterning activity of sonic hedgehog by secreted BMP inhibitors expressed by notochord and somites. Development 127, 4855–4866.

44. Liu, J., Merkle, F.T., Gandhi, A.V., Gagnon, J.A., Woods, I.G., Chiu, C.N., Shimogori, T., Schier, A.F., and Prober, D.A. (2015). Evolutionarily conserved regulation of hypocretin neuron specification by Lhx9. Development 142, 1113–1124.

45. Loo, L., Simon, J.M., Xing, L., McCoy, E.S., Niehaus, J.K., Guo, J., Anton, E.S., and Zylka, M.J. (2019). Single-cell transcriptomic analysis of mouse neocortical development. Nat. Commun. 10, 134.

46. Lu, F., Kar, D., Gruenig, N., Zhang, Z.W., Cousins, N., Rodgers, H.M., Swindell, E.C., Jamrich, M., Schuurmans, C., Mathers, P.H., et al. (2013). Rax is a selector gene for mediobasal hypothalamic cell types. J. Neurosci. 33, 259–272.

47. Manning, L., Ohyama, K., Saeger, B., Hatano, O., Wilson, S.A., Logan, M., and Placzek, M. (2006). Regional morphogenesis in the hypothalamus: a BMP-Tbx2 pathway coordinates fate and proliferation through Shh downregulation. Dev. Cell 11, 873–885.

48. Marion, J.-F., Yang, C., Caqueret, A., Boucher, F., and Michaud, J.L. (2005). Sim1 and Sim2 are required for the correct targeting of mammillary body axons. Development 132, 5527–5537.

49. Mathieu, J., Barth, A., Rosa, F.M., Wilson, S.W., and Peyriéras, N. (2002). Distinct and cooperative roles for Nodal and Hedgehog signals during hypothalamic development. Development 129, 3055–3065.

50. Mayer, C., Hafemeister, C., Bandler, R.C., Machold, R., Batista Brito, R., Jaglin, X., Allaway, K., Butler, A., Fishell, G., and Satija, R. (2018). Developmental diversification of cortical inhibitory interneurons. Nature 555, 457–462.

51. Melsted, P., Booeshaghi, A.S., Liu, L., Gao, F., Lu, L., Min, K.H.J., da Veiga Beltrame, E., Hjörleifsson, K.E., Gehring, J., and Pachter, L. (2021). Modular, efficient and constant-memory single-cell RNA-seq preprocessing. Nat. Biotechnol.

52. Merkle, F.T., Maroof, A., Wataya, T., Sasai, Y., Studer, L., Eggan, K., and Schier, A.F. (2015). Generation of neuropeptidergic hypothalamic neurons from human pluripotent stem cells. Development 142, 633–643.

53. Müller, F., Albert, S., Blader, P., Fischer, N., Hallonet, M., and Strähle, U. (2000). Direct action of the nodal-related signal cyclops in induction of sonic hedgehog in the ventral midline of the CNS. Development 127, 3889–3897.

54. Nagasaki, H., Kodani, Y., and Suga, H. (2015). Induction of Hypothalamic Neurons from Pluripotent Stem Cells. Interdisciplinary Information Sciences 21, 261–266.

55. Newman, E.A., Wu, D., Taketo, M.M., Zhang, J., and Blackshaw, S. (2018). Canonical Wnt signaling regulates patterning, differentiation and nucleogenesis in mouse hypothalamus and prethalamus. Dev. Biol. 442, 236–248.

56. Ohyama, K., Ellis, P., Kimura, S., and Placzek, M. (2005). Directed differentiation of neural cells to hypothalamic dopaminergic neurons. Development 132, 5185–5197.

57. Padilla, S.L., Carmody, J.S., and Zeltser, L.M. (2010). Pomc-expressing progenitors give rise to antagonistic neuronal populations in hypothalamic feeding circuits. Nat. Med. 16, 403–405.

58. Patten, I., Kulesa, P., Shen, M.M., Fraser, S., and Placzek, M. (2003). Distinct modes of floor plate induction in the chick embryo. Development 130, 4809–4821.

59. Pearson, C.A., Ohyama, K., Manning, L., Aghamohammadzadeh, S., Sang, H., and Placzek, M. (2011). FGF-dependent midline-derived progenitor cells in hypothalamic infundibular development. Development 138, 2613–2624.

60. Placzek, M., and Briscoe, J. (2005). The floor plate: multiple cells, multiple signals. Nature Reviews Neuroscience 6, 230–240.

61. Romanov, R.A., Tretiakov, E.O., Kastriti, M.E., Zupancic, M., Häring, M., Korchynska, S., Popadin, K., Benevento, M., Rebernik, P., Lallemend, F., et al. (2020). Molecular design of hypothalamus development. Nature 582, 246–252.

62. Salvatierra, J., Lee, D.A., Zibetti, C., Duran-Moreno, M., Yoo, S., Newman, E.A., Wang, H., Bedont, J.L., de Melo, J., Miranda-Angulo, A.L., et al. (2014). The LIM homeodomain factor Lhx2 is required for hypothalamic tanycyte specification and differentiation. J. Neurosci. 34, 16809–16820.

63. Saper, C.B., and Lowell, B.B. (2014). The hypothalamus. Curr. Biol. 24, R1111–R1116.

64. Seifinejad, A., Li, S., Mikhail, C., Vassalli, A., Pradervand, S., Arribat, Y., Pezeshgi Modarres, H., Allen, B., John, R.M., Amati, F., et al. (2019). Molecular codes and in vitro generation of hypocretin and melanin concentrating hormone neurons. Proc. Natl. Acad. Sci. U. S. A. 116, 17061–17070.

65. Shimogori, T., Lee, D.A., Miranda-Angulo, A., Yang, Y., Wang, H., Jiang, L., Yoshida, A.C., Kataoka, A., Mashiko, H., Avetisyan, M., et al. (2010). A genomic atlas of mouse hypothalamic development. Nat. Neurosci. 13, 767–775.

66. Staudt, N., and Houart, C. (2007). The prethalamus is established during gastrulation and influences diencephalic regionalization. PLoS Biol. 5, e69.

67. Swaab, D.F. (2003). Human Hypothalamus: Basic and Clinical Aspects, Part I (Elsevier).

68. Trapnell, C., Cacchiarelli, D., Grimsby, J., Pokharel, P., Li, S., Morse, M., Lennon, N.J., Livak, K.J., Mikkelsen, T.S., and Rinn, J.L. (2014). The dynamics and regulators of cell fate decisions are revealed by pseudotemporal ordering of single cells. Nat. Biotechnol. 32, 381–386.

69. Wang, L., Egli, D., and Leibel, R.L. (2016). Efficient Generation of Hypothalamic Neurons from Human Pluripotent Stem Cells. Current Protocols in Human Genetics 90.

70. Ware, M., Hamdi-Rozé, H., and Dupé, V. (2014). Notch signaling and proneural genes work together to control the neural building blocks for the initial scaffold in the hypothalamus. Front. Neuroanat. 8, 140.

71. Ware, M., Hamdi-Rozé, H., Le Friec, J., David, V., and Dupé, V. (2016). Regulation of downstream neuronal genes by proneural transcription factors during initial neurogenesis in the vertebrate brain. Neural Dev. 11, 22.

72. Wolf, F.A., Angerer, P., and Theis, F.J. (2018). SCANPY: large-scale single-cell gene expression data analysis. Genome Biol. 19, 15.

73. Xie, Y., and Dorsky, R.I. (2017). Development of the hypothalamus: conservation, modification and innovation. Development 144, 1588–1599.

74. Xu, S., Yang, H., Menon, V., Lemire, A.L., Wang, L., Henry, F.E., Turaga, S.C., and Sternson, S.M. (2020). Behavioral state coding by molecularly defined paraventricular hypothalamic cell type ensembles. Science 370, eabb2494.

75. Yoo, S., Kim, J., Lyu, P., Hoang, T.V., Ma, A., Trinh, V., Dai, W., Jiang, L., Leavey, P., Won, J.-K., et al. (2021). Control of neurogenic competence in mammalian hypothalamic tanycytes.

76. Zhou, X., Zhong, S., Peng, H., Liu, J., Ding, W., Sun, L., Ma, Q., Liu, Z., Chen, R., Wu, Q., et al. (2020). Cellular and molecular properties of neural progenitors in the developing mammalian hypothalamus. Nat. Commun. 11, 4063.

